# Mechanically coupled proliferation controls notochord geometry during axis elongation

**DOI:** 10.64898/2026.11.13.765900

**Authors:** Kasumi Kishi, Shi-Lei Xue, Marco Dalla Vecchia, Edouard Hannezo, Anna Kicheva

**Author notes:** equal contribution (Kasumi Kishi, Shi-Lei Xue).

## Abstract

The notochord is a central organizing structure in vertebrate development, functioning as a signaling axis that patterns surrounding tissues while undergoing rapid elongation. Despite its importance, how notochord shape is regulated to ensure robust morphogen signaling remains poorly understood. Here, we combine high-resolution 3D imaging of mouse embryos between E8.5 to E10.5 of development with a data-driven biophysical model to investigate the mechanisms underlying notochord morphogenesis. We find that the notochord maintains a remarkably constant cross-sectional area along most of its anterior–posterior extent, a feature that is preserved over time and in mutants with reduced body axis length. To uncover the cellular basis of this geometric robustness, we quantified cell behaviours and their contribution to notochord morphogenesis. We show that notochord expansion is driven by a posterior-to-anterior gradient in cell proliferation, accompanied by changes in cell size and directed posterior cell migration. These processes act in a coordinated and complementary manner to offset the stress gradient generated by axis extension. In particular, our analysis indicates that the proliferation gradient is regulated by mechanical feedback from surrounding tissues, ensuring the maintenance of a constant trunk cross-sectional area throughout elongation. Together, these findings show that organ morphogenesis is tightly coordinated with the surrounding and provide a framework for understanding how tissue shape is robustly controlled in growing tissues.

## Introduction

As growing organs are shaped during development, local growth processes must be coordinated with mechanical forces generated intrinsically as well as by neighboring tissues. This coordination is particularly challenging during body axis elongation, where multiple tissues alter their shape while remaining physically associated with one another (Tam and Behringer, 1997). Mechanical feedback has emerged as a key regulator of tissue growth and morphogenesis across tissues (Heisenberg and Bellaïche, 2013; Mongera et al., 2019), yet how individual organs sense and respond to the mechanical environment to maintain appropriate geometry during development is poorly understood.

The notochord is a midline mesodermal structure that plays a central role during trunk development in chordates. In birds, amphibians, and fish, the notochord contributes structurally to axis elongation, and in all vertebrates it functions as a key signaling center for surrounding tissues (Stemple, 2005). The notochord secretes morphogens that regulate pattern formation and tissue growth, influencing the adjacent neural tube and paraxial mesoderm. In the neural tube, Sonic hedgehog (Shh) produced by the notochord is essential for dorsoventral patterning and for the specification of the floor plate (Placzek and Briscoe, 2005). Perturbations of notochordal Shh expression or notochord integrity lead to pronounced defects in neural patterning, underscoring the sensitivity of this system to notochord-derived signals (Ho et al., 2024; Yu et al., 2013). Since Shh acts in a dose-dependent manner, the size of the notochord is likely a key determinant of patterning outcomes — yet how notochord size is controlled during mouse development remains unclear.

In mouse embryos, the notochord develops at gastrulation from the node and notochordal plate, both of which express T/Brachury (Beddington, 1994; Kinder et al., 2001; Sulik et al., 1994; Ukita et al., 2009; Wilkinson et al., 1990). As the trunk extends, the notochord undergoes substantial elongation. Tissue elongation can result from multiple cellular processes, including cell proliferation, rearrangements, and changes in cell size or shape. During their residence in the ventral node between E7.5 and E8.25, notochord precursor cells exhibit little proliferative activity, with only ∼1% of cells incorporating BrdU (Bellomo et al., 1996; Ukita et al., 2009). Following this period, at E9.5 and E10.5, cells in the posterior notochord display active proliferation (Ukita et al., 2009). Cell divisions and the rearrangement of cells from the node into the posterior notochord have been shown to contribute to rostrocaudal elongation of the mouse notochord (Sausedo and Schoenwolf, 1994). Yet, how these cellular and mechanical processes are integrated to result in robust anisotropic morphogenesis during notochord elongation is poorly understood.

In addition to tissue-intrinsic cell behaviours, growing tissues such as the notochord are subjected to mechanical stresses that arise from their physical coupling to neighboring tissues. At the early stages of notochord formation, mechanical pressure from the amniotic sac regulates notochord elongation (Imuta et al., 2014). As the notochord transitions to a rod-like structure, it builds up an enveloping extracellular matrix (ECM) layer, containing collagens, proteoglycans and glycoproteins (Trapani et al., 2017). Mice with perturbed notochord ECM have aberrant notochord shapes (Guo et al., 2020; Smits and Lefebvre, 2003). At the same time, the posterior notochord is further coupled to the surrounding presomitic mesoderm (PSM), which has been suggested to compress axial tissues – thus ablation of the PSM in chick embryos impairs axial tissue elongation (Xiong et al., 2020). Later in development, the ECM sheath of the zebrafish notochord is mechanically linked to the adjacent somite boundaries and this is required for correct notochord segmentation (Wopat et al., 2024). Altogether, these observations indicate that notochord shape reflects a combination of intrinsic cell behaviours, ECM mechanics and forces from surrounding tissues. Yet the quantitative contribution of each of these factors, as well as whether and how they feedback onto each other to encode the final morphogenetic shape, remains to be determined.

These gaps highlight the need for a quantitative, multiscale analysis of notochord growth in the mouse embryo that links cellular behaviors, tissue mechanics, and organ-level coordination during body axis elongation. To address this, we quantified the size and shape of the mouse notochord between E8.5 and E10.5 of development. We found that despite substantial lengthening, the notochord maintains a stable cross-sectional area in the trunk – which is not a priori expected and implies active control of tissue geometry. By systematically characterizing notochord cell behaviours, we found that notochord extension occurs primarily through cell proliferation. By establishing a biophysical model of notochord growth and morphogenesis in combination with genetic perturbations, we found that control of cell proliferation by mechanosensitive feedback can explain the maintenance of a well-defined cross-sectional area while allowing the notochord length to match the body axis extension. Our results establish a principle by which mechanical coupling between a growing organ and its environment provides the spatial information required to tune cell proliferation and maintain anisotropic organ geometry during morphogenesis.

## Results

### The notochord lengthens as the body extends and keeps its trunk cross-sectional area constant

To obtain precise quantitative measurements of the size and shape of the developing mouse notochord, we developed a stage-specific tissue clearing protocol and combined it with immunostaining against T/Brachury (hereafter referred to as Bra) to mark notochord cells. This approach allowed us to visualize the entire organ in whole embryos between E8.5 and E10.5 of development (7 and 39 somite stages, respectively) (Fig. 1A, Fig. S1A). In order to reconstruct the notochord shape from 3D z-stacks of whole embryos, we developed a custom image analysis pipeline (Methods) and validated it by comparing to tissue cryosections (Fig. 1B, C and S1B-D). Notochords are first straightened along the AP axis, then individual nuclei are segmented and a volume mask is created, allowing us to measure the volume, length and cross-sectional area of the notochord along its length (Fig. 1C, S1B).

**Figure 1:**
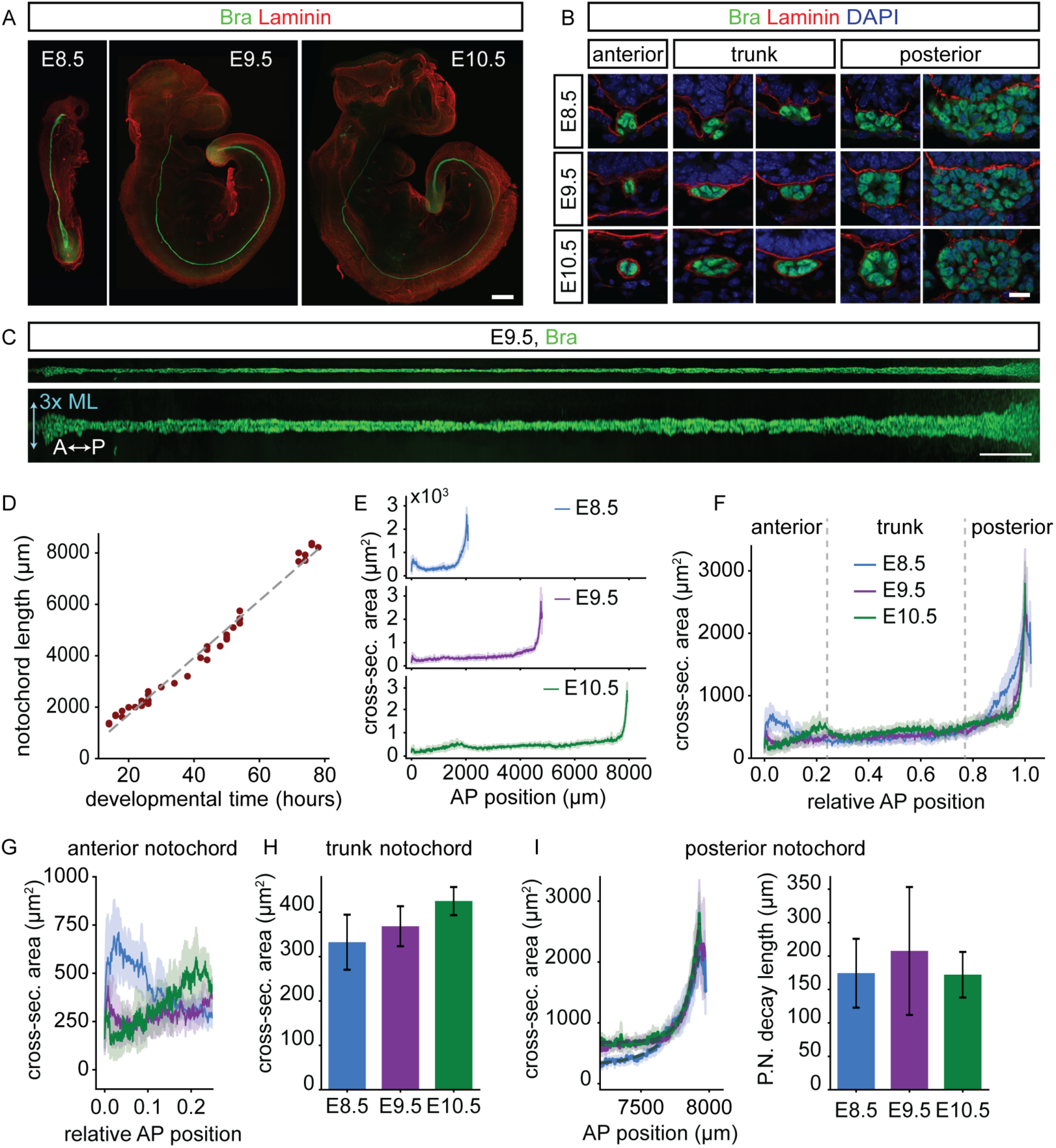
(A) Maximum projections of cleared mouse embryos from E8.5 to E10.5 immunostained against Bra and Laminin. Scale bar 300µm. (B) Notochord cross-sections organized by relative AP position at indicated stages. Immunostaining against Bra and Laminin. Scale bar 15µm. (C) Digitally straightened E9.5 notochord, at original scale (upper image) and 3x mediolaterally enlarged (lower image). Scale bar 200µm. (D) Quantification of notochord length from E8.5 to E10.5 (dots). Embryo stage (h) was estimated from the somite stage x 2h/somite. Linear fit (dashed) corresponds to *l*(*t*) = *L*_0_ + *u_tip_*(*t* − *t*_0_) with *t*_0_ = 18ℎ, *L*_0_ ≈ 1485*μm*, *u_tip_* ≈ 111*μm*/ℎ. (E) Mean notochord cross-sectional area along the AP axis from E8.5 to E10.5 (Methods). Shaded regions, 95% CI. n= 12 for E8.5, n=9 for E9.5, n=6 for E10.5. (F) Mean notochord cross-sectional area and 95% CI as a function of relative AP position. Vertical dotted lines demarcate three regions as defined in the main text. (G) Magnified view of the anterior region in F. (H) Mean area for the whole trunk region at the indicated stages. Error bars, SD. (I) Mean notochord cross-sectional area in the posterior region. Profiles are registered to the E10.5 profile, such that the posterior peak position = 7927µm. Exponential fits to the AP area decay (dashed, E8.5 green, E9.5 pink/purple, E10.5 blue) with estimated decay length (right panel).

Our data indicated that the notochord length increased approximately 4-fold between E8.5 and E10.5 of development, from ∼2000 µm to 8000 µm, which corresponds to an increase of 2664 µm/day or 111 µm/h (Fig. 1D). Quantification of the cross-sectional area revealed that it varies along the AP axis, delineating three distinct regions (Fig. 1E-I). The anterior region corresponds to the most anterior 25% of the AP length and spans the level of the head and cervical somites, up to somite 4 at E10.5 (Fig. 1F). The trunk region encompasses the middle portion of the AP axis, spanning 25% to 80% of its length, while the posterior region covers the most posterior 20% and includes the unsegmented zone and after E8.5, the posterior-most somites. In the anterior region, the cross-sectional area profile varied over time and the mean decreased approximately 2-fold from E8.5 to E10.5 (Fig. 1F, G). By contrast, the cross-sectional area in the trunk region was remarkably constant along the AP axis (Fig. 1F) and on average increased only ∼1.3-fold over developmental time (Fig. 1H). The cross-sectional area in the posterior region was significantly higher than the rest of the notochord. It reached a maximal value close to the posterior end that was ∼7-fold higher than the mean area in the trunk region, and decreased exponentially towards the anterior (Fig. 1F, I). Notably, both the amplitude and decay length of this exponential decline remained approximately constant over time (Fig. 1I), indicating that the notochord has a near-constant funnel-like shape.

Together, these data indicate that the notochord elongates substantially as the body axis extends while keeping its trunk cross-sectional area approximately constant. These observations raise the question of how the trunk notochord area is determined and maintained.

### Cell divisions account for the observed notochord growth

To investigate the cell behaviors that underlie the observed changes in notochord shape between E8.5 and E10.5 of development, we systematically analyzed cell size, cell rearrangements, cell proliferation and cell death. Quantifications of the cell density (Methods) indicated that it is constant along the AP axis and increases approximately ∼1.2 fold between E8.5 and E10.5 of development (Fig. 2A). This indicates that cell size becomes smaller over time (Fig. S2A), hence the length of the notochord does not increase because of increasing cell size. Instead, this suggests that the notochord size increases by an increase in cell number.

**Figure 2:**
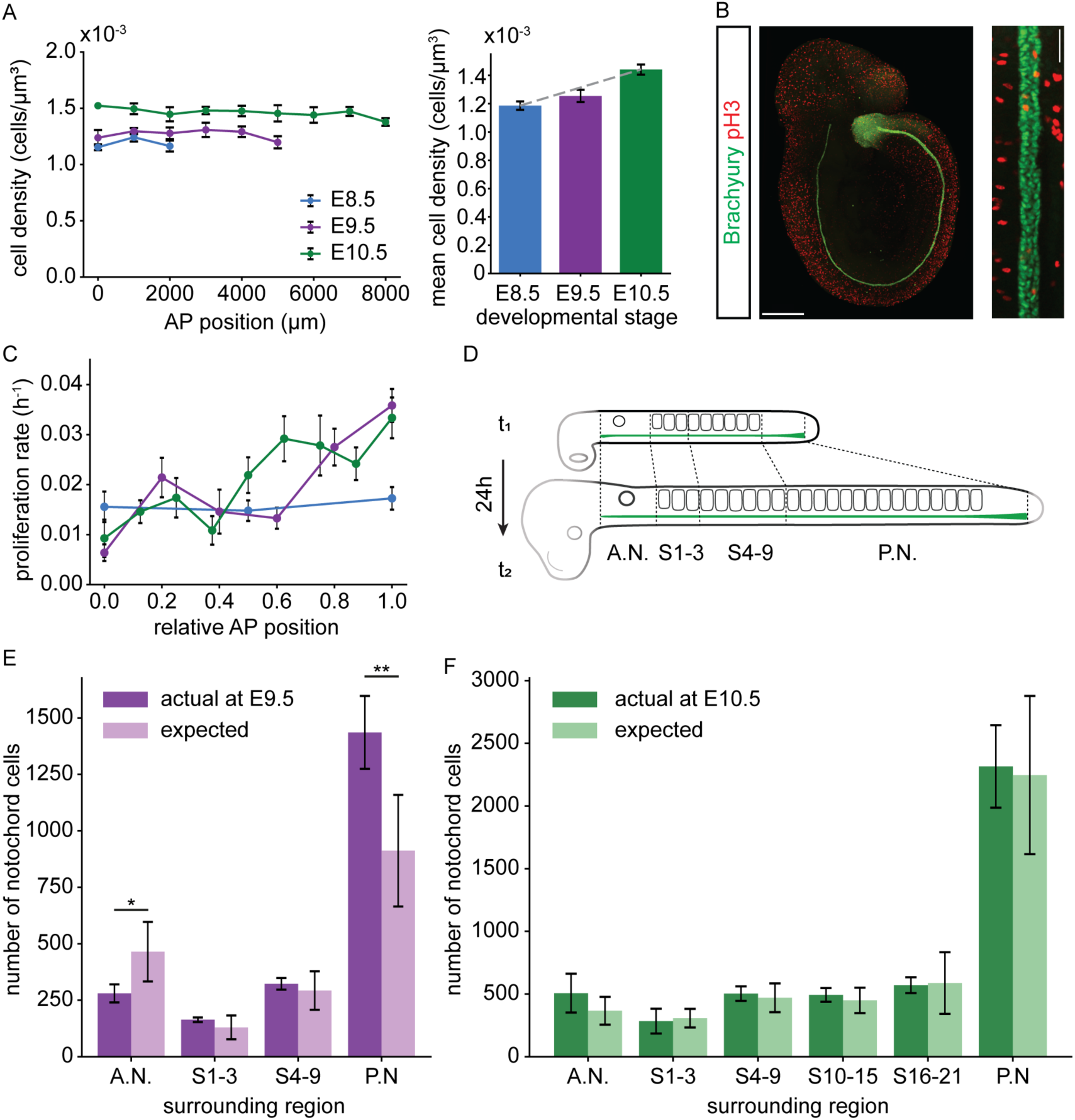
(A) Quantification of notochord cell density along the AP axis at the indicated stages. Mean cell density for the whole notochord (right panel). Grey dotted line shows the fit for *ρ* = *ρ*_0_(1 + *k_ρ_t*) where *k_ρ_* = 0.0045ℎ^−1^. Error bars, SD. Sample size E8.5 n=12, E9.5 n=9, E10.5 n=6. (B) Left panel, E9.5 mouse embryo immunostained against Brachyury and pH3. Scale bar 400µm. Right panel, zoomed region after digital straightening, scale bar 50µm. (C) Proliferation rate along the AP axis at the indicated stages estimated from pH3-mitotic fraction (methods). Error bars, S.E.M. Sample size E8.5 n=12, E9.5 n=9, E10.5 n=6. (D) Schematic illustrating the region landmarks used for the quantification in E. (E-F) Measured number of cells at E9.5 (E) and E10.5 (F) and number of cells expected from the measured proliferation rate (C) and cell number at E8.5 (E) and E9.5 (F), respectively. Error bars, SD. Two-sample t-test p-value >0.05 unless otherwise indicated, * p-value=0.033, ** p-value=0.006.

To analyze the increase in cell number, we next investigated the rate of cell division. We subdivided the notochord into regions of ∼1000 µm in AP length and inferred the proliferation rate from the fraction of cells in mitosis using immunostainings against phospho-Histone 3 and Bra (Fig. 2B). These data indicated that at E8.5 the proliferation rate was nearly constant along the AP axis and corresponded to ∼0.016 h^-1^, or a cell cycle length of ∼43 h (Fig. 2C). This is consistent with previous reports (Bellomo et al., 1996; Ukita et al., 2009) showing that the notochord division rate is very low at these developmental stages. By contrast, at E9.5 and E10.5, the proliferation rate had a graded spatial profile that increased from ∼0.008 h^-1^ in the most anterior regions to ∼0.035 h^-1^ (corresponding to cell cycle length of ∼18 h) at the posterior (Fig. 2C). This suggests that the notochord cell number increases over time primarily by proliferation of the fast-dividing posterior region at later stages. Inspection of the level of apoptosis using cleaved caspase 3 immunostaining showed that cell loss is overall negligible, except in the most anterior region at E8.5 and E9.5 (Fig. S2B, C). These data further support the conclusion that the number of cells in the notochord increases primarily through fast cell proliferation in the posterior region from E9.5 onwards.

To assess whether the proliferation rate alone is sufficient to account for the observed increase in number of notochord cells -or whether additional cells are added from a Bra-low stem cell pool -we compared the measured total cell numbers at E9.5 and E10.5 to those predicted from the measured proliferation rate (Fig. 2C) and notochord cell number 24h earlier, at E8.5 and E9.5, respectively. To do this, we subdivided the notochord into discrete regions along the AP axis (Fig. 2D). This analysis revealed very good correspondence between measured and predicted cell counts throughout the AP axis (Fig. 2E, F), with the exception of the anterior region at E9.5, where the actual cell count was ∼200 cells lower than expected, and the most posterior region at E9.5, in which there were ∼500 cells more than expected (Fig. 2E). This suggests that cell proliferation of Bra-positive notochord cells is sufficient to account for notochord growth after E9.5, but there is a small contribution of posterior cell accretion prior to E9.5.

Together, these data indicate that notochord growth is the highest in the posterior region, which is characterized by a large cross-sectional area, higher proliferation, lower apoptosis, and posterior cell accretion prior to E9.5. This suggests a tight and regulated balance of mechanical forces, in order to remodel the newly generated cells at the thick posterior region into an invariant and thin shape of the trunk notochord. We therefore asked whether cell rearrangements and directed movements contribute to distributing cells along the anteroposterior axis during elongation.

### Notochord cells undergo fluid, coherent posterior migration

To investigate the rearrangements of notochord cells during morphogenesis, we established a protocol for live imaging of mouse trunk explants for up to 5 h (Fig. 3A, Methods). To this end, we dissected and cultured explants containing the trunk and posterior regions of E9.5 embryos expressing Shh-CreERT2-inducible H2B-mCherry or membrane EGFP. We first quantified the orientation of daughter cells during cell division using H2B-mCherry to identify telophase spindles (Fig. 3B). This quantification revealed that daughter cells tend to align along the AP axis at the end of mitosis (Fig. 3C). This raised the possibility that oriented cell divisions may facilitate the unidirectional AP growth of the notochord if the relative positions of daughter cells are maintained without further rearrangements. However, tracking the relative distance of daughter cells over time revealed that it increased linearly, reaching a characteristic cell diameter (∼10 µm) after ∼ 5h (Fig. 3D, E, Supplementary Movie M1), indicating that cells continuously drift apart. Consistent with this, in 30% of the tracked cell pairs we observed one or more neighbor exchanges (defined as events in which the distance between the cells exceeded 10µm) (Fig. S2D). Thus, the tissue is fluid on the timescales of morphogenesis (days) and short-term oriented cell divisions are unlikely to translate to long-term directional elongation.

**Figure 3:**
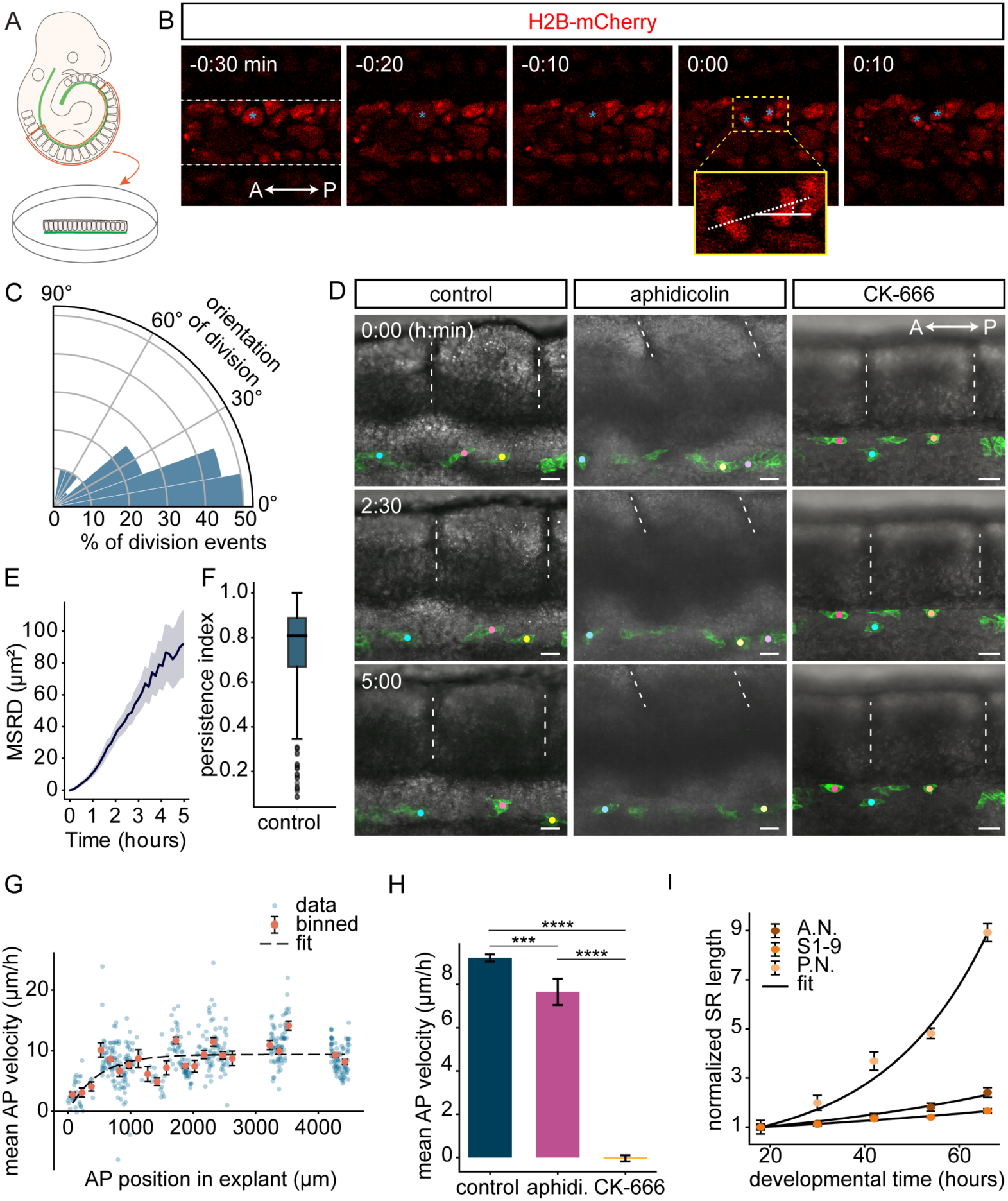
**(A)** Schematic of E9.5 trunk explant preparation for live imaging. **(B)** Time-lapse snapshots showing a cell division (asterisks) in the notochord (white outline). Inset (yellow) shows division angle (dotted line) relative to the AP axis (solid white line). Posterior, right. **(C)** Cell division orientation relative to the AP axis (0°). Sample size, n=36. KS-test p-value<0.0001 indicates non-random distribution. **(D)** Time-lapse imaging of notochord sparsely induced with membrane GFP in the presence of DMSO (control), 40µM aphidicolin or 800µM CK-666. Colored circles, tracked cells. Dashed lines, somite boundaries. Time stamps rounded to nearest half hour. Scale bar 40µm. Posterior, right. **(E)** Mean squared relative distance (MSRD) between tracked neighbouring cells over time. Sample size, n=272 cell pairs. **(F)** Persistence index of tracked notochord cells (Methods). Sample size, n=470 cells. **(G)** Quantification of AP notochord cell velocity (Methods, anterior explant boundary = 0 µm). Blue points, mean velocity of individual cells. Orange points, mean velocity of cells in 150µm-length bins ± SEM. The length scale *l_c_* was estimated by fitting the binned data (dashed line, Methods). **(H)** Mean AP cell velocity across the explant for control, aphidicolin-and CK-666-treated explants (concentrations as in D). Sample size control n=470, aphidicolin n=85, CK-666 n=130. One way ANOVA, p<0.0001; asterisks p-values from Tukey’s HSD post hoc test, *** p=0.0007, **** p<0.0001. **(I)** AP length of regions surrounding the notochord over time with exponential fit (solid line). A.N. region (anterior to first somite), S1-9=somites 1 to 9, P.N. somite 10 to posterior notochord edge. Length is measured at the midline and normalized to the initial time point (E8.5).

Strikingly, tracking also revealed that notochord cells displayed pronounced and persistent displacement in posterior direction with respect to the nearest adjacent somite boundary (Fig. 3D, F). Quantification of this movement demonstrated that, except for the most anterior ∼500 µm, cells move with a constant mean velocity of 9.2±0.2 µm/h (mean ± SEM), equivalent to ∼ 221 µm per day, relative to somite boundaries throughout the AP length of the explant (Fig. 3G). Cells close to the anterior edge of explants appeared to move more slowly (∼2.8 µm/h) with respect to adjacent somite boundaries, suggesting that cells require a certain distance to reach maximum velocity. The net posterior orientation of cell movement raised the possibility that the movement is driven by active cell migration, rather than passive displacement generated by cell divisions. To test whether this is the case, we cultured mouse explants in media containing 40µM aphidicolin to inhibit cell division (Fig. S2E) or 800µM of the Arp2/3 inhibitor CK-666, which blocks actin-mediated active cell migration (Krndija et al., 2019), but leaves cell divisions intact in the time course of our experiment (Fig. S2F). We then tracked the movement of cells in trunk positions and compared them to DMSO-treated controls (Fig. 3D, Supplementary movies M2, M3). Remarkably, while the cell velocity in aphidicolin-treated explants was maintained to levels similar to control (83 % of control value), cell displacement was abrogated in CK-666-treated explants (Fig. 3H). This indicates that the observed posterior cell displacement relative to somite boundaries in explants results from active cell migration, rather than passive displacement due to cell divisions.

The observation that cells in the trunk region appear stationary with respect to somite boundaries in CK-666-treated explants further implies that in the absence of active cell migration, the notochord and the surrounding somites extend in register, without significantly ‘sliding’ with respect to each other. In the presence of active cell migration, there is posterior displacement of the trunk notochord corresponding to 221 µm per day. This posterior sliding of the notochord with respect to the surrounding body generated by active cell migration is approximately 12-fold lower in magnitude compared to the overall extension generated by cell division, which corresponds to ∼2.7 mm per day (Fig. 1D). Altogether, the small magnitude of the cell migration relative to the overall extension and the considerable rearrangements of cells suggest that the main factor underlying notochord morphogenesis is the gradient of the rate of cell proliferation, while cell migration has a minor contribution.

### A biophysical model captures the observed changes in notochord shape

Our findings so far revealed that the notochord elongates primarily through a cell proliferation gradient that peaks at the posterior end. While the length of the notochord continuously increases to match the surrounding body, the cross-sectional area is maintained largely constant. This suggests that the rate of cell division must be tuned such that it generates the required AP extension, while at the same time, additional constraints ensure that notochord growth does not occur radially. To investigate potential regulatory mechanisms, we developed a minimal biophysical model of notochord morphogenesis.

In this model, the notochord is represented as a viscous axisymmetric rod with cross-sectional area *A(x, t)* varying along the AP axis *x* and over time. The evolution of notochord shape is governed by a conservation equation:

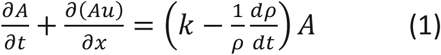

where the change in cross-sectional area *A* in time and space depends on the velocity *u*(*x*, *t*) of notochord cells relative to the anterior boundary and on the notochord growth rate, which incorporates the proliferation rate *k* and changes in cell density *ρ*(*t*). Cell density was observed to increase linearly with time as *ρ* = *ρ*_0_(1 + *k_ρ_t*), where the rate constant *k_ρ_* = 0.0045/ℎ was determined from the quantification in Fig. 2A. The anterior and posterior boundaries of the notochord are identical to those of the surrounding tissue and correspond to *u*(*x* = 0) = 0 at the anterior and *u*(*x* = *l*) = d*l*/dt at the posterior end, where *l*(*t*) is the notochord length.

Furthermore, the notochord shape can be described in terms of the mechanical forces that act on it. Notochord cells move with an active migration force *f_m_*, have a tissue viscosity of *η* and experience a friction force *f* = *ξ*(*u* − *u_e_*) with respect to the surrounding tissue, where *u_e_* is the velocity of the external tissue relative to the anterior boundary and *ξ* is the friction coefficient. Therefore, the force balance equation is:

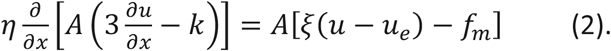

We first sought to estimate key parameters based on experimental data. To understand how the notochord moves relative to the surrounding, we first considered the growth kinetics of the paraxial mesoderm surrounding the notochord. To this end, we quantified the distances between defined somite positions and morphological landmarks along the AP axis as illustrated in Fig. 2D at successive developmental stages (Fig. 3I), from which we computed the local AP extension rates (Methods). These data revealed that the paraxial mesoderm extends with an approximately 3-4-fold slower AP extension rate in the head and cervical regions (0.011-0.018 h^-1^), compared to the trunk together with tail region (0.045 h^-1^) (Fig. 3I). Thus, similar to the notochord, the surrounding body undergoes faster extension posteriorly.

To incorporate this observation into the model, we approximated the experimentally observed posterior-to anterior decline of the extension velocity of external tissues *u_e_*(*x*, *t*) at each developmental stage (Fig. 4A, Methods) as an exponential function 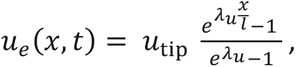 where *λ_u_* (*t*) is the characteristic spatial decay length and *u*_tip_ is the velocity of the posterior tip (Fig. 4B, Methods). To estimate *u*_tip_, we considered that starting from an initial length *L*_0_, the body elongates with velocity *x* resulting in total length *l*(*t*) = *L*_0_ + *u*_tip_*t*. A linear fit to the measured notochord length over time (Fig. 1D) allowed us to estimate *L*_0_ =1485µm at somite stage 9 and *u*_tip_ = 111 *μm*/ℎ. With this estimate of *u*_tip_, we inferred *λ_u_*(*t*) by fitting *u_e_*(*x*, *t*) (Fig. 4A, B).

**Figure 4:**
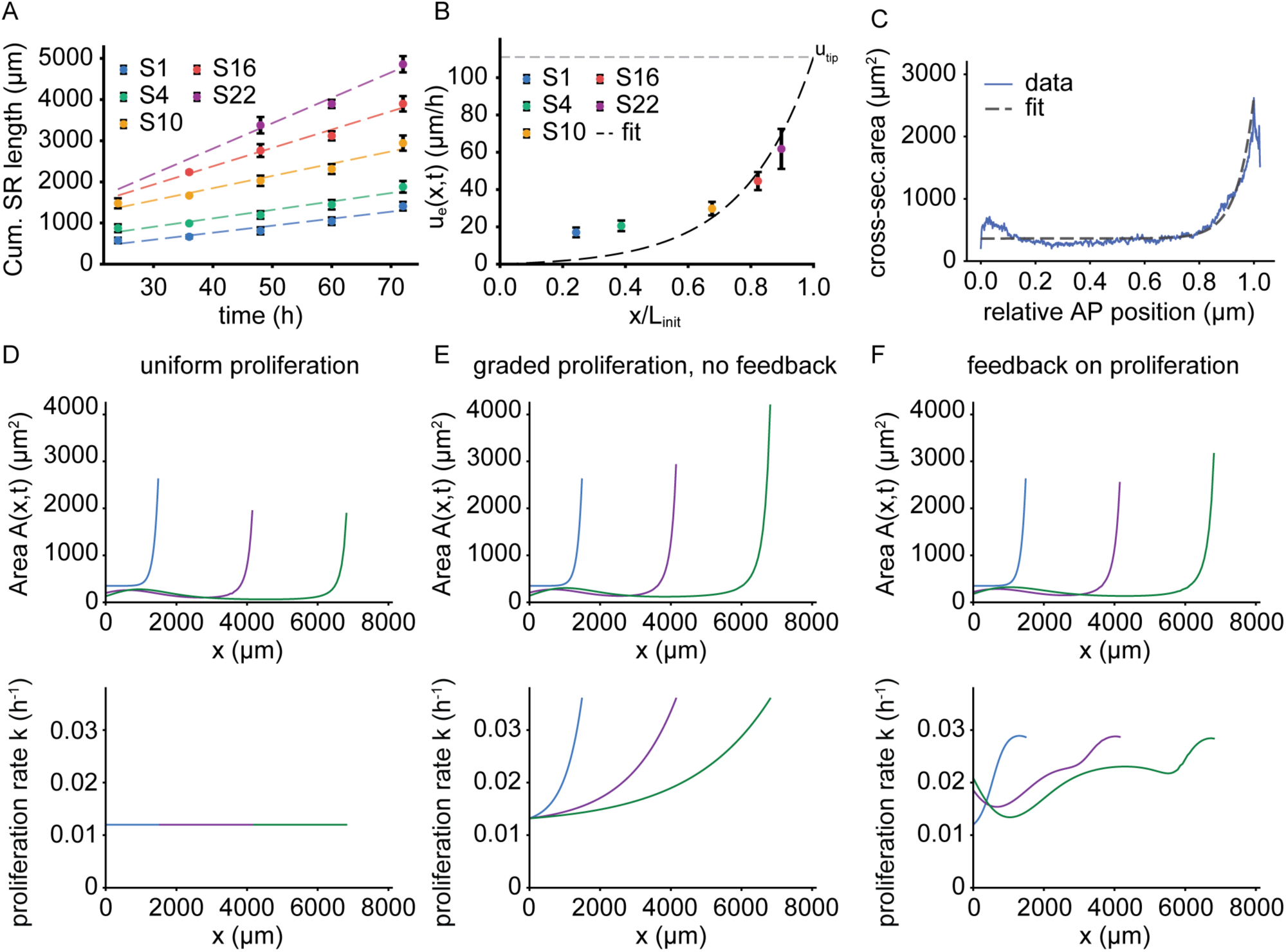
**(A)** Cumulative length of surrounding tissues over time measured at the midline from the notochord anterior edge to the indicated somite landmarks. Error bars, SD. Dashed lines, linear fits *Cum*. *SR* = *Cum*. *SR_tinit_* + *u_e_*(*x*, *t*) ⋅ *t*. **(B)** Slopes from linear fits in (A) (color-coded as in A) against relative AP position of the somite landmark at the initial time point. *L_init_* = 2028*μm* is the mean notochord length at E8.5. Black curve, fit to *u_e_*(*x*, *t*) with *λ_u_* = 4.5 and *u_tip_* = 111*μm*/ℎ (Methods). Error bars, ±1 SE of fit in (A). **(C)** Quantification of the mean notochord cross-sectional area at E8.5 and the fitted equation for *A*(*x*, 0) (Methods). **(D)** Simulated notochord cross-sectional area (top) for a constant proliferation rate across AP position and time (bottom). **(E)** Simulated notochord cross-sectional area (top) for an exponential proliferation rate profile along the AP axis (bottom). **(F)** Simulated notochord cross-sectional area (top) and cell proliferation (bottom) where the cell proliferation rate is regulated through feedback.

Non-dimensionalization of the model yields a characteristic length scale 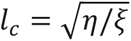 which relates the magnitude of the friction force to the viscous force. This parameter represents the distance needed for migrating cells to accelerate from a stationary state (at the anterior boundary) to their maximum migration speed in the trunk (Methods). To determine the approximate magnitude of this parameter, we fit the model to the measured cell velocities in mouse explant experiments (Fig. 3G, Methods). This revealed that a characteristic length of 275 µm describes the observed increase in speed from 2.8 µm/h close to the anterior boundary to a constant migration velocity of 9.4 µm/h in the trunk region.

The remaining parameter to predict the evolution of the cross-section area is the growth rate *k*. To understand its impact, we varied *k* and assessed the area profiles over time, using an exponential fit to the measured area profile at E8.5 as an initial condition *A*_0_(*x*) (Fig. 4C). Spatially and temporally uniform *k* (k=0.012 h^-1^), of the same order as the experimentally measured proliferation rate (Fig. 2C), failed to maintain the trunk and peak posterior cross-sectional areas over time and led to progressive notochord thinning due to body elongation (Fig. 4D). By contrast, a growth rate that increases towards the posterior (Methods) yields area profiles with a stable trunk region and a sustained posterior peak (Fig. 4E), consistent with experimental observations (Fig. 1H, I). This analysis shows that the spatiotemporal profile of the proliferation rate *k* critically determines the evolution of *A*(*x*, *t*). This raises the question of how proliferation is regulated to achieve a robust notochord shape.

### Notochord proliferation is coupled to axis elongation

In order to investigate the mechanisms that control notochord shape, we considered a scenario in which posterior extension is perturbed. In Loop-tail (LPT) mouse mutants, a defect in the PCP gene Vangl2 results in a shortened AP axis (Greene et al., 1998; Ybot-Gonzalez et al., 2007). Consistent with this, we observed that homozygous LPT mutant embryos have smaller somite and notochord lengths compared to control littermates from E9.5 onwards, and reach ∼80% of the control length at E10.5 (Fig. 5A, B, S3A). A linear fit to the notochord length over time (Fig. 5B) indicated that the tip extension velocity *u_tip_* decreased to ∼71% of its wildtype value and corresponded to 79 µm/h, while *L*_0_ remained similar. As expected from smaller external stresses on the notochord (Fig. S3B, C), reducing AP embryo extension resulted in a larger posterior notochord area (Fig. 5C). Our data indicated that the amplitude of the posterior area was up to 2.7-fold higher in LPT mutants (Fig. 5C, S3D). In contrast to the posterior area, LPT embryos at all developmental stages have trunk notochord areas comparable to those of control littermates (Fig. 5C, S3E).

**Figure 5:**
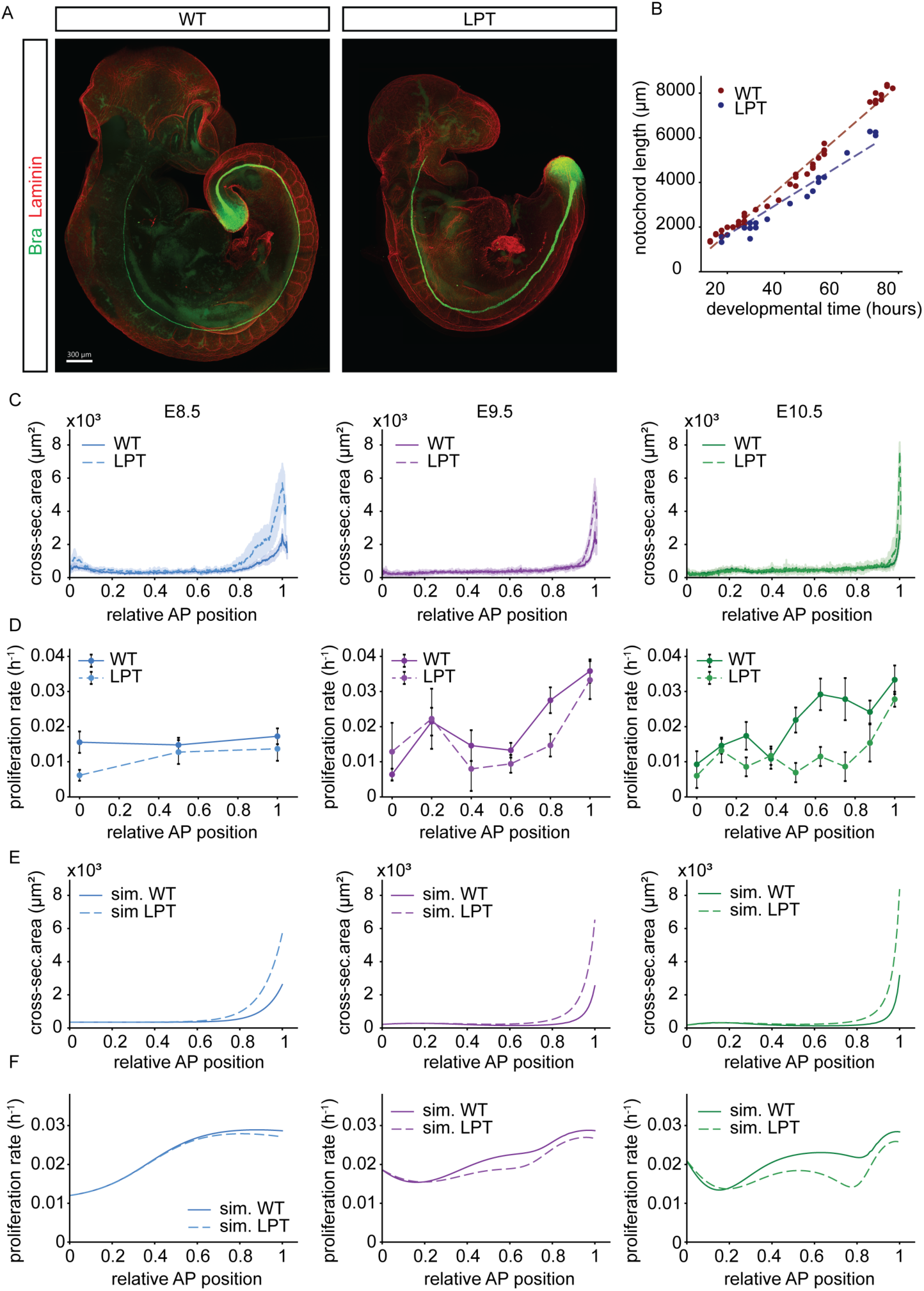
**(A)** E9.5 wildtype and homozygous LPT embryos immunostained against Bra. **(B)** Notochord length of wildtype (red, same as in Fig. 1D) and LPT embryos (blue) between E8.5 and E10.5 of development. Linear fit (dashed) *l*(*t*) = *L*_0_ + *u_tip_*(*t* − *t*_0_) yields *u_tip_* ≈ 79*μm*/ℎ for LPT. **(C)** Notochord cross-sectional area profiles of wildtype (solid) and homozygous LPT (dashed) embryos between E8.5 and E10.5. Shaded regions, 95% confidence intervals. Sample size E8.5 n=6, E9.5 n=5, E10.5 n=4. **(D)** Quantifications of the notochord proliferation rate (based on pH3 staining, Methods) in wildtype and LPT embryos between E8.5 and E10.5. Error bars, SEM. **(E)** Simulated notochord cross-sectional area for control and LPT conditions (Methods) for the model with feedback. **(F)** Simulations of the notochord cell proliferation rate, conditions as in E.

These observations show that the trunk area is robust to changes in the body elongation velocity and raise the question of whether the growth rate in the trunk changes in the LPT mutants to compensate for the decreased elongation. To test this, we measured the proliferation rate in LPT mutants using pH3 immunostaining. Strikingly, this showed that the proliferation rate decreased compared to control embryos at all stages from E8.5 to E10.5 (Fig. 5D), while the cell density remained unchanged (Fig. S3F). This suggests that the proliferation rate is modulated to maintain a constant trunk cross-sectional area, a feature not included in the current model.

In order to understand how the growth rate is tuned to generate the correct area profiles, we investigated possible theoretical constraints on the growth rate. In particular, we considered a mechanism whereby a default growth rate in the absence of feedback *k*_0_ is modulated by the deviation of the area from a specific “threshold” area *A_e_* with strength *m*: 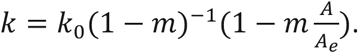 Similar mechanical feedback control of proliferation has been previously considered in other systems (Alessandri et al., 2013; Basan et al., 2009; Dolega et al., 2021; Ranft et al., 2010; Shraiman, 2005) and could be compatible with multiple molecular mechanisms, for instance, confinement of notochord cells by stiff extracellular matrix. To capture this position-dependence, we defined 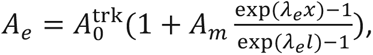 where *A_m_* and *λ_e_* are unknown parameters that tune the spatial profile of the feedback (Methods). Numerical simulations of this model indicated that it reliably captures the experimentally observed area profiles in wildtype embryos (Fig. 4F), similar to the model without feedback (Fig. 4E). To test the behavior of the model in LPT mutants, we simulated the area profiles for reduced u_tip_ (u_tip_=79µm, corresponding to the experimental measurement (Fig. 5B)). The simulation results (Fig. 5E, F) showed that as expected, the proliferation rate in the trunk region of LPT mutants decreased compared to controls (Fig. 5F). Crucially, the model with feedback can reliably capture the experimentally observed area profiles in LPT mutants: the trunk area was maintained while the posterior peak area increased ∼2.6-fold (Fig. 5E). Overall, although both models, with or without feedback, could explain the increased area of the posterior notochord in LPT mutants, the presence of a feedback mechanism on growth enables the trunk notochord area to be less sensitive to external changes in elongation (Fig. S3G).

To more directly validate the feedback mechanism, we asked how the notochord responds to perturbations that reduce its area, which the model predicts should lead to an increase in the proliferation rate (Fig. 6A, B). To test this, we generated embryos (hereafter referred to as DTA) in which tamoxifen-inducible Shh-CreERT2 transiently and mosaically induces recombination that allows diphtheria toxin subunit A expression in notochord cells, thereby causing apoptosis (Fig. 6C, S4A). Upon tamoxifen injection at E6.5, we observed cleaved caspase 3 positive notochord cells at E8.5 (Fig. 6D). Subsequently, at E9.5 and at E10.5, parts of the notochord were missing (Fig. 6C, E). A closer inspection indicated that the basement membrane, marked by Laminin immunostaining, disassembled and disappeared in the missing regions, while the remaining notochord fragments retained their ECM, but their cross-sectional area decreased at E9.5 and E10.5 (Fig. 6F, G, S4B). Consistent with this phenotype, we observed defects in DV patterning of the overlying neural tube that correlated in severity with corresponding notochord defect (Fig. S4C). Crucially, the overall proliferation rate in the mutant embryos measured with pH3 immunostaining was increased compared to controls (Fig. 6H-I) at E9.5, consistent with the model predictions. Consistent with feedback from the notochord cross-sectional area, rather than response to local tissue damage or cell density, mitotic cells were uniformly distributed along fragments and the cell density within fragments was similar between DTA and control conditions (Fig. S4D, E). At E10.5, the proliferation rate in DTA mutants was similar to controls (Fig. 6H), although the area had not fully recovered, which suggests that the feedback is transient or imperfect. Nevertheless, our E9.5 data support the existence of feedback between the notochord area and the rate of proliferation.

**Figure 6:**
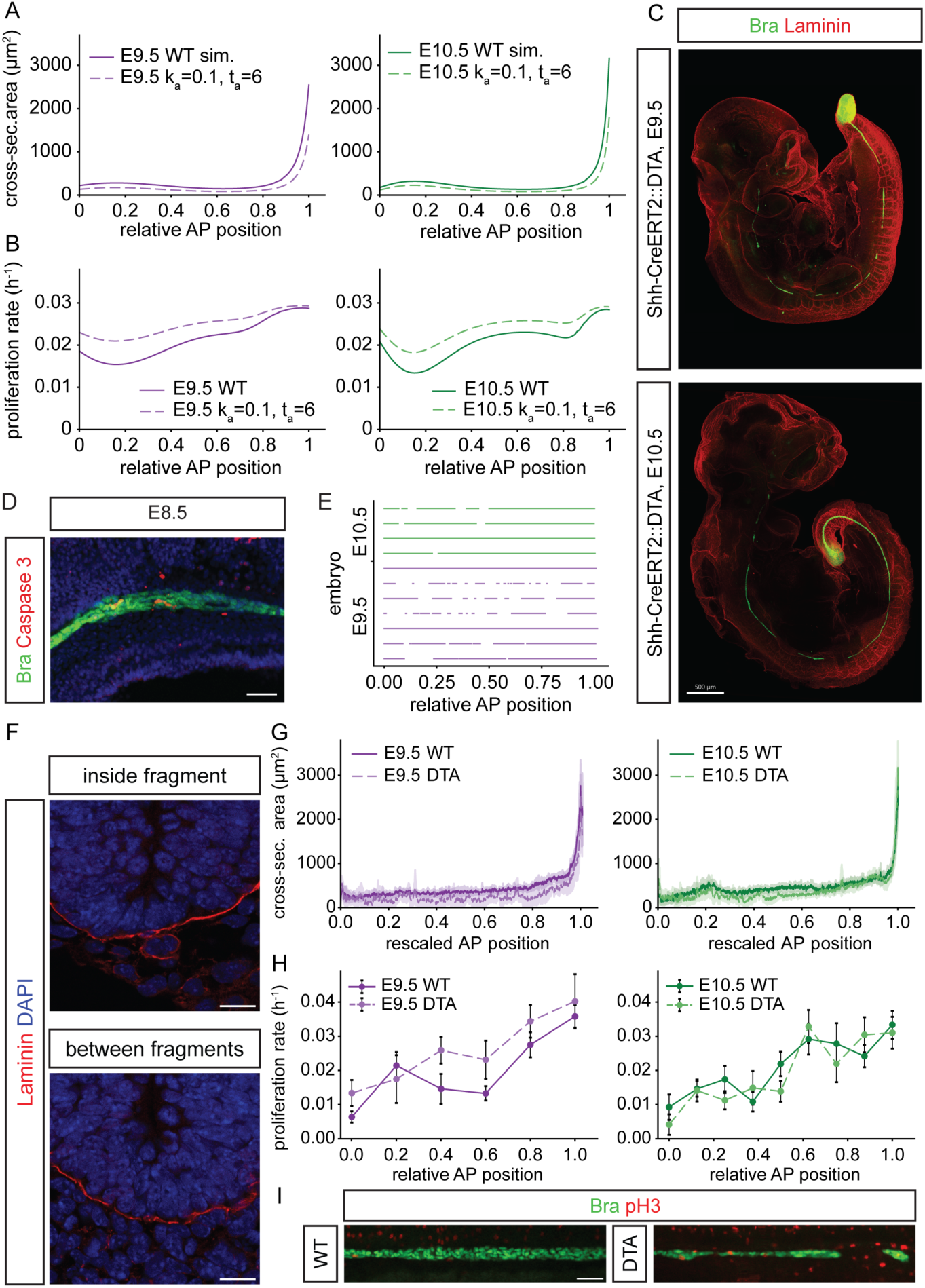
**(A)** Simulated notochord cross-sectional area for wildtype vs DTA condition, implemented by inducing cell death for ta=6h at a rate of k_a_=0.1h^-1^ from the initial time point t=18 (E8.5). **(B)** Predictions of the notochord cell proliferation rate in control and DTA conditions from simulations, as in (A). **(C)** E9.5 and E10.5 DTA embryos immunostained against Bra injected with tamoxifen at E6.5. Scale bar 500µm. **(D)** DTA-induced notochord cell death at E8.5 visualized by immunostaining against cleaved caspase 3. Scale bar 50 µm. **(E)** Locations of notochord fragments (line) and gaps in DTA embryos at E9.5 and E10.5, injected with tamoxifen at E6.5. **(F)** Image of E9.5 DTA embryo cross-sections showing the presence and absence of Laminin surrounding the notochord in fragments and in gap regions, respectively. Scale bar 15µm. **(G)** Quantification of mean notochord cross-sectional area in wildtype and DTA embryos at E9.5 and E10.5. Shaded region, 95% confidence interval. Sample size E8.5 n=6, E10.5 n=8. **(H)** Quantifications of the notochord cell proliferation rate in WT and DTA embryos at E9.5 and E10.5. Error bars SEM. Sample size E8.5 n=6, E10.5 n=8. **(I)** Immunostaining against phospho-Histone 3 (pH3) and Bra shows mitotic cells in the notochord of WT and DTA embryos. Scale bar, 50µm.

### The active migration of notochord cells counterbalances elongation-induced tissue thinning

Body axis extension generates a stress gradient that is maximal in the posterior region of the notochord and tends to decrease the notochord area posteriorly. Our analysis suggests that this is counterbalanced by the proliferation rate gradient. Next, we asked whether the active posterior cell migration that we observed (Fig. 3D-H) may have a similar role in compensating for the local notochord thinning that body axis extension tends to create. To this end, we performed simulations in which we progressively reduced the active cell migration velocity *u_m_*. Reducing the active migration velocity leads to a concomitant decrease in the area of the notochord at the posterior tip (Fig. 7A). Analysis of the area profile upon reduction of *u_m_* by 20% of its default value in our simulations indicated that the trunk area is maintained constant in this condition, while the anterior area increases (Fig. 7B). Thus, despite the relatively small magnitude of active migration speed, the model suggests that it plays a role in tuning the cross-sectional area in specific regions.

**Figure 7:**
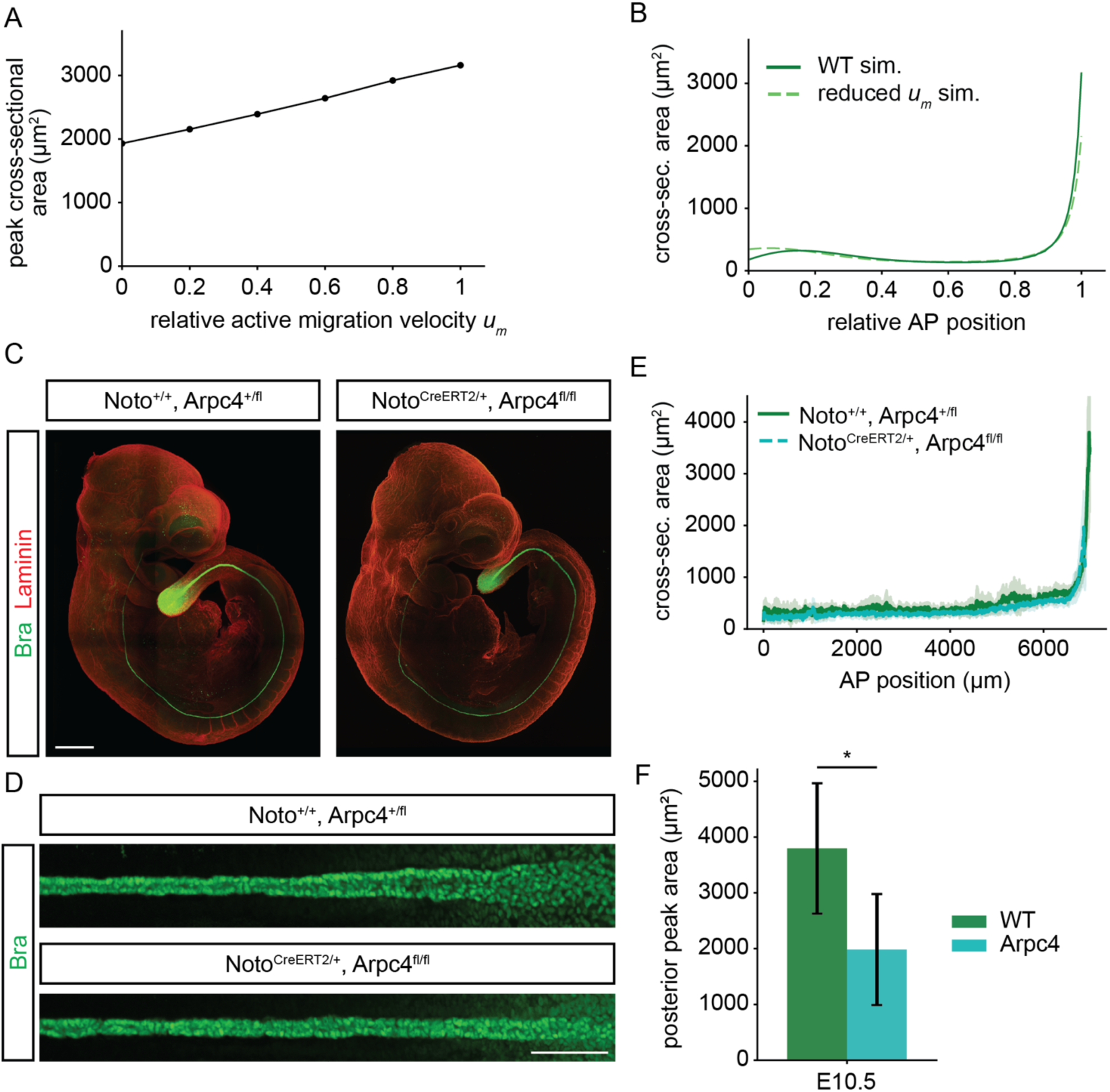
**(A)** Relationship between the peak posterior cross-sectional area and the relative active migration velocity *u_m_* in simulations. **(B)** Predicted notochord cross-sectional area at E10.5 when the active migration velocity is set to 20% of control levels in simulations. **(C)** E10.5 control and Noto^CreERT2/+^::Arpc4^fl/fl^ embryos, immunostained against Bra and Laminin. Scale bar 500 µm. **(D)** Images of the posterior notochord of E10.5 control and Noto^CreERT2/+^::Arpc4^fl/fl^ embryos after digital straightening. Scale bar 100µm **(E)** Quantification of the notochord cross-sectional area in E10.5 control and Noto^CreERT2/+^::Arpc4^fl/fl^ embryos. Shaded regions, 95% CI. Sample size n=5 for control, n=9 for Noto^CreERT2/+^::Arpc4^fl/fl^. **(F)** Mean posterior peak cross-sectional area in E10.5 control and Noto^CreERT2/+^::Arpc4^fl/fl^ embryos. Sample size as in E. Two-sided independent samples t-test, p-value=0.032.

To experimentally test the predicted effects of cell migration, we generated mouse embryos in which the Arpc4 subunit of the Arp2/3 complex was deleted by Noto-CreERT2 mediated recombination by tamoxifen injection at E6.5. This mutation was reported to abrogate active actin-dependent cell migration in other systems (Krndija et al., 2019). Consistent with the model predictions, we observed reduced posterior notochord cross-sectional area in these mutants at E10.5, while the trunk area was maintained constant (Fig. 7C-F). The mean anterior area was not changed, although it is possible that the Cre-induced variability in the mutant condition precludes the detection of a phenotype in the anterior region in our assay. Altogether, these results suggest that the active posterior migration of notochord cells counterbalances the high stress generated by body axis elongation.

## Discussion

Here, we combined quantitative imaging with biophysical modeling to show that the notochord trunk cross-sectional area is actively maintained during vertebrate body axis elongation, through feedback regulation of the proliferation rate. Our results support a model in which body axis elongation generates a posterior stress gradient that tends to thin the notochord, and this is counterbalanced by a gradient in proliferation and directed cell movement, ensuring robust control of notochord cross-sectional area. Crucially, we show that the proliferation rate is actively regulated by feedback to maintain a robust cross-sectional area despite the mechanical demands of elongation.

A key feature of our model is that the elongation of the surrounding tissues is explicitly coupled to the notochord length. This coupling arises naturally from the fact that notochord, the surrounding PSM and neural tube all arise through a dynamic process of cell fate specification in the node and caudo-lateral epiblast during gastrulation (Cambray and Wilson, 2007; Tzouanacou et al., 2009), sharing a posterior growth zone. In addition, the notochord and PSM are physically connected at the tailbud via cell-to fibronectin matrix adhesion (Dray et al., 2013) – loss of this connection in integrin zebrafish morphants (itga5mo;aVmo) causes the tissues in the tailbud to detach and the notochord to undulate. Besides having a common posterior endpoint with the external tissues, the notochord experiences friction with respect to these tissues along its AP length.

Our live imaging data in mouse explants showed that the notochord and somites do not undergo AP extension in register. Instead, there is active posterior cell migration of notochord cells relative to the adjacent somites in the trunk, suggesting that the notochord extends more slowly than the adjacent tissues in its posterior region. Consistent with this, the elongation rate of the PSM and posterior somites (0.045/h) is higher than the proliferation rate of the notochord in this region (0.035/h), with the difference accommodated by posterior migration.

Previous work in zebrafish has demonstrated that notochord elongation is dependent on the formation of fluid-filled vacuoles that lead to the enlargement of notochord cells over time (Ellis et al., 2013; McLaren and Steventon, 2021). Vacuolization of the notochord is required for correct axis extension (Ellis et al., 2013), consistent with the crucial structural support role of the notochord in this species. While large fluid-filled vacuoles are characteristic of notochord cells in fish, amphibians and birds (Stemple, 2005), these are not observed in the mouse notochord. Instead, our measurements indicate that mouse notochord cells decrease in volume over time, thus increasing the cell density (Fig. 2A). Mouse notochord elongation is therefore driven primarily by cell proliferation rather than vacuole-mediated expansion. Interestingly, the earlier anterior vacuolization of notochord cells in zebrafish results in posterior displacement of notochord cells relative to the adjacent tissues (McLaren and Steventon, 2021). Similar to these findings, we also observed posterior displacement of notochord cells relative to the adjacent somites in mouse embryos. However, our study indicates that in mouse this is driven by active cell migration, rather than passive displacement due to cell enlargement (Fig. 3D, H). These findings suggest that posterior displacement of the notochord cells relative to the surrounding tissues is a conserved feature of vertebrate axis elongation, achieved by distinct cellular mechanisms in different species (vacuole-driven expansion in zebrafish and active migration in mouse).

A key finding of our study is that the proliferation rate in the mouse notochord adjusts to generate the required length without altering the trunk cross-sectional area. We show that this is achieved in two ways. First, the maximum posterior proliferation rate is maintained regardless of body elongation, suggesting that newly generated notochord cells have intrinsically faster cell cycles (as found in many developing tissues), or are in an external signaling environment that maintains fast proliferation. Second, the trunk proliferation rate is modulated to compensate for deviations in the cross-sectional area, as evidenced by the response to transient apoptosis and slowed elongation. How this feedback is molecularly and mechanically controlled is currently unknown. A plausible mediator of this coupling is the extracellular matrix, which has been reported to link axial tissues and transmit mechanical signals (Bénazéraf et al., 2010; Dray et al., 2013; Guillon et al., 2020; Wopat et al., 2024). It is conceivable that embryo elongation could modulate the mechanical properties of the ECM surrounding the notochord, for example its stiffness or organization, which in turn feeds back on cell proliferation. In this view, ECM stiffness may depend on the time elapsed since its deposition, providing a potential mechanism by which the feedback is spatially patterned resulting in constraining the trunk but not the posterior notochord cross-sectional area. While the underlying pathways remain to be defined, this type of feedback is effectively captured by our target area model, in which growth is regulated in response to the evolving mechanical state of the tissue.

The robustness of notochord cross-sectional area has direct implications for Shh gradient formation in the tissues adjacent to the notochord. Neural tube and paraxial mesoderm are first exposed to Shh emanating from the posterior notochord region and the levels of this early Shh exposure are key for floor plate formation in the neural tube (Ho et al., 2024; Placzek and Briscoe, 2005; Yu et al., 2013). Consistent with the increased posterior notochord area in LPT mutants, LPT embryos appear to have expanded floor plate domains (Greene et al., 1998). Furthermore, our DTA experiment demonstrates that local aberrations in notochord cross-sectional area lead to corresponding local patterning defects in the overlying neural tube (Fig. S4C). This suggests that the robustness of notochord trunk cross-sectional area contributes to a robust and continuous DV pattern formation along the AP axis in adjacent tissues. Despite the constant area in the notochord trunk, the overall amplitude of the Shh gradient in the neural tube increases over time (Cohen et al., 2015). Our data raises the possibility that this results from the temporal increase in cell density. Alternatively, the production rate per cell or the degradation dynamics change over time. Future studies would be needed to understand how morphogenesis affects the Shh gradient dynamics.

Together, our findings reveal that mouse notochord elongation is achieved through a dynamically regulated interplay between tissue deformation and growth. By uncovering a feedback between elongation and proliferation that preserves tissue geometry, our work highlights how local cell behaviors are tuned to global morphogenetic demands. These results suggest that coordination between the notochord and its surrounding environment is essential for robust axis extension, and provide a framework for understanding how developing tissues achieve precise control of shape through mechanically mediated growth regulation.

## Acknowledgements

We thank Danijela Matic Vignjevic for the Arpc4^fl^ mice and the ISTA IOF and PCF SSUs. AK and KK are supported by ISTA, the European Research Council under Horizon Europe: grant 101044579 and Austrian Science Fund (FWF): F78 (Stem Cell Modulation). SX is supported by the National Natural Science Foundation of China (Grant No. 12402073). EH is supported by ISTA.

## Author Contributions

KK conceptualization, investigation, methodology, formal analysis, software, validation, visualization, writing – review and editing. SX conceptualization, formal analysis, software, visualization, writing – review and editing. MDV methodology, software. EH conceptualization, supervision, writing – review and editing. AK conceptualization, supervision, writing – original draft, writing – review and editing.

## Methods

### Mouse lines

All work with animals was approved under the license BMWFW-66.018/0006-WF/V/3b/2016 from the Austrian Bundesministerium für Wissenschaft, Forschung und Wirtschaft. Аll procedures were performed in accordance with the relevant regulations. Mouse lines were maintained on a CD-1 background unless specified otherwise. The following transgenic lines used in the study were previously described: ShhCreERT2 (JAX:005623, (Harfe et al., 2004) NotoCreERT2 (Imuta et al., 2013), mT/mG (JAX:007676, (Muzumdar et al., 2007)), Rosa26-LSL-H2B-mCherry, abbreviated H2B-mCherry (JAX:023139, (Peron et al., 2015)), Vangl2LP, also known as Loop tail (LPT) (JAX:000220, (Murdoch, 2001)), ROSA-DTA (JAX: 009669,(Voehringer et al., 2008)), and Arpc4fl (Krndija et al., 2019).

For embryo collection and analysis, timed matings were set up in the afternoon and checked for a vaginal plug the following morning. The following timed matings with tamoxifen-inducible Cre were used: ShhCreERT2/+ crossed to ROSA-DTA to generate ShhCreERT2/+, Rosa-DTA/+ embryos referred to as DTA throughout the manuscript; NotoCreERT2/+, Arpc4fl/+ crossed to Arpc4fl/+ to generate NotoCreERT2/+, Arpc4fl/fl embryos. To induce temporally controlled Cre expression, tamoxifen was injected in pregnant mothers at the following time points and doses: 4mg tamoxifen injected at E6.5 for ROSA-DTA; 3mg at E6.5 for Arpc4fl/fl.

For mouse explant live imaging experiments, ShhCreERT2/+ was crossed to mT/mG and pregnant mothers were injected with 1.7 mg tamoxifen at E6.5. Embryos that expressed membrane EGFP sparsely in the notochord were selected and collected at E9.5. Some of the experiments were performed with mT/mG females maintained on a 129/SV background instead of a CD-1 background and injected with 1.4mg tamoxifen - this did not result in any phenotypic difference to CD-1 females. Both backgrounds were used for the quantifications shown. Live imaging experiments were also performed with H2B-mCherry to track the cell nuclei. In these experiments, ShhCreERT2/+ was crossed to H2B-mCherry, 3mg tamoxifen was injected at E6.5, and embryos were collected at E9.5.

### Live imaging and analysis of mouse trunk explants

#### Sample preparation and imaging

Embryos were dissected in media prewarmed to 37°C, consisting of 90% DMEM/F12 (Gibco, 21041025), 10% FBS (Gibco, 10270106), and 1X pen/strep (Gibco, 15140122). Tissues anterior to somite 4 and ventral to the notochord were removed. Tissues posterior to somite 21 were also removed in some experiments – these behaved identical to explants with posterior tip intact, hence both types of explants were used for the quantifications shown. The explant was oriented ventral down in a 35mm glass bottom dish and stabilized in a drop of freshly made solution of 8mg/ml fibrinogen (Merck, 341573) and 0.5U/ml thrombin (Sigma, T7326) (Januschke and Gonzalez, 2010; Kicheva et al., 2014). Subsequently, 1.2ml media consisting of 40% DMEM/F12, 50% rat serum (prepared in-house), 10% FBS and 1X pen/strep was added. Explants were imaged at a ZEISS inverted LSM800 confocal microscope, using a Plan APOCHROMAT 20x objective (ZEISS, 420650-9901). Z-stacks were obtained at 0.9μm intervals for ShhCreERT2, mT/mG explants and at 1μm intervals for ShhCreERT2, H2B-mCherry explants. Images were acquired every 9-15 minutes over the course of 6 hours. Samples were kept at 37°C, 5% CO2 and 90% humidity throughout live imaging.

#### Analysis of cell movement and rearrangements

For analysis of cell migration, images were imported to Fiji, rotated to align the AP axis of the explant to the horizontal X axis of the image, and Z-drift was corrected using the plugin “Correct 3D drift” (Parslow et al., 2014). Notochord cells and the positions of somite boundaries were manually tracked in Imaris. A rolling mean was applied to smooth the tracked positions with a rolling window of 3 time steps. The displacement of the somite boundaries in the X (AP) axis were subtracted from the displacement of notochord cells, to calculate the relative displacement of notochord cells to somites. The mean AP notochord cell velocity was estimated as 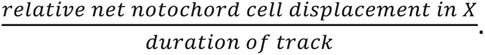

To calculate the persistence index, the total displacement of the cell (the distance between the cell at the first and last time point of the track) was divided by the total path length (the sum of the distance travelled between each time point), for every cell track.

To quantify neighbour exchanges, cell pairs were classified as neighbours if the distance between their centroids was less than 10 µm. A neighbour exchange event was defined as the time point at which this inter-centroid distance increased by more than 10 µm relative to its value at the start of the tracking window, indicating loss of contact. Subsequent exchange events were identified when the distance changed by more than 10 µm relative to the distance recorded at the preceding exchange event, capturing both loss and re-establishment of contact.

The mean squared relative displacement (MSRD) was quantified for pairs of cells as defined above. At each time point, the distance between the two cells was calculated from their smoothed coordinates. Then, the MSRD was calculated at every time point as *MRSD* = (*distance*(*t*) − *distance*(*t*_0_))T^2^.

#### Cell division

Aphidicolin (Sigma, A0781) and CK-666 (Sigma, SML0006) were dissolved in DMSO to make 10mM and 100mM stock, respectively. To determine the effect of aphidicolin on cell proliferation, explants were prepared as described above, and cultured for 6 hours at 37°C, 5% CO2 and 90% humidity at the indicated final concentrations. Subsequently, explants were fixed in 4% PFA for 1 hour, immunostained against T/Bra and pH3 as described in Table 4, and imaged at a ZEISS inverted LSM800 confocal microscope. The mitotic index was calculated as the number of pH3+ T+ double-positive cells divided by the notochord length in the explant. The division rate of notochord cells in the presence of CK-666 was determined by counting the number of mitosis events during the live imaging experiment. The length of the notochord in the observation region was measured, and the division rate was calculated as 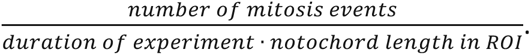

To analyze the orientation of cell division, mitotic cells were tracked using H2B-mCherry. The angle between the axis of division and the AP axis of the explant was measured at telophase. Angle measurements were performed in Fiji (Schindelin et al., 2012). The acute angle was taken ([0°, 90°]), such that 0° and 90° correspond to a division angle parallel or orthogonal to the AP axis respectively.

### Whole embryo clearing

Embryos were fixed on ice in 4% PFA for the following durations: E8.5 for 1.25 hours, E9.5 for 1.5 hours, and E10.5 for 2 hours. Embryos were washed three times in PBS post-fixation, once in PBS + 0.1% TWEEN (PBSTween), and stored at 4°C for up to two days. To perform tissue clearing, the protocol from (Pende et al., 2020) was modified as follows. Embryos underwent dehydration in increasing concentrations of methanol (25%, 50%, 75%, 100%) in PBSTween. Embryos were stored in 100% methanol at 4°C overnight, or at -20°C until further processing. E8.5 and E9.5 embryos were rehydrated in decreasing methanol concentrations (75%, 50%, 25%, 0%) in PBSTween. E10.5 embryos were incubated in a 1:2 methanol:H2O2 solution overnight before undergoing serial rehydration. Embryos were cleared at 37°C at 250rpm using the clearing solution and duration in Table M1. After clearing, embryos were washed 5 times in PBSTween at room temperature and used directly for immunohistochemistry.

**Table M1.** Clearing solution composition and clearing duration for different embryonic stages.

| Developmental stage | Clearing solution (in water) | Clearing duration |
|---|---|---|
| **E8.5** | 5% Triton X-100<br>8% THEED (Sigma, 87600)<br>5% Urea | 2 minutes |
| **E9.5** | 5% Triton X-100<br>10% THEED<br>5% Urea | 6 minutes |
| **E10.5** | 5% Triton X-100<br>8% THEED<br>10% Urea<br>1% CHAPS | 13 minutes |

### Immunohistochemistry and microscopy of cleared embryos

All steps were performed at 4°C. Cleared embryos were incubated with primary antibodies in blocking buffer overnight, and subsequently subjected to 4 short washes (x 5 min each in PBSTween) and a long wash in PBSTween. This was followed by incubation with secondary antibodies and DAPI in blocking buffer, 4 short washes, 1 long wash, and mounting for imaging. Durations of primary and secondary antibody incubations and long washes, as well as blocking buffer composition and antibody concentrations were varied according to developmental stage as given in Table M2. Primary and secondary antibodies used are described in Table M3. Embryos were mounted on glass slides in EasyIndex mounting medium (RI=1.46, Life Canvas Technologies, DLC/EI-Z101).

**Table M2.** Blocking buffer, antibody dilutions and incubation durations used for whole embryo immunostaining at different embryonic stages.

| Embryonic stage | Blocking buffer (in PBS) | Primary antibodies | Secondary antibodies | Duration of antibody incubations and long washes |
|---|---|---|---|---|
| **E8.5-E9.5** | 0.2% Tween<br>1% BSA | Goat T (1:80)<br>Rat pH3 (1:800)<br>Rabbit Laminin (1:800)<br>Rabbit CC3 (1:200) | anti goat FITC (1:150)<br>anti rat pH3 (1:800)<br>anti rabbit Alexa Fluor 647 (1:150)<br>500µg/ml DAPI (1:50) | overnight |
| **E10.5** | 0.3% Triton X-100<br>1% BSA<br>1% CHAPS<br>1% 2,6-Di-O-methyl-β-cyclodextrin (TCI Chemicals, D5992) | Goat T (1:50)<br>Rat pH3 (1:600)<br>Rabbit Laminin (1:600)<br>Rabbit CC3 (1:100) | Anti goat FITC (1:50)<br>anti rat pH3 (1:600)<br>anti rabbit Alexa Fluor 647 (1:50)<br>500µg/ml DAPI (1:50) | 2 days |

Cleared embryos were imaged using a Nikon CSU-W1 Spinning Disk microscope using a CFI Plan Apochromat 10x/0.45 air (MRD00105) objective. Z-stacks encompassing the embryo were imaged with a Z-interval of 2.5μm, and 12-bit tile scan images were acquired with 5% overlap. Alternatively, cleared embryos were imaged using an Andor Dragonfly 505 microscope with the spinning disk modality using a CFI Plan Apochromat 10x/0.5 glycerol objective (MRD71120). Z stacks were imaged with an interval of 2.5μm, and 16-bit tile scan images were acquired with 5% overlap.

### Analysis of notochord shape and cell number dynamics

Images of cleared embryos were converted from ND2 to Imaris format using the Imaris File Converter (version 9.9.1, Bitplane), stitched using the Imaris Stitcher (version 9.9.1, Bitplane), and exported to tif using the Bio-Formats importer (Linkert et al., 2010) in Fiji (Schindelin et al., 2012). A digitally straightened image of the notochord was generated in Fiji by manually tracing the Bra+ notochord along its AP length in XY and XZ maximum intensity projections using the segmented line tool. The built-in “straighten” tool was used to straighten the image along the manually traced line in both axes.

To distinguish the notochord signal from background, two independent models were trained using the pixel classification workflow in Ilastik (Berg et al., 2019). The nuclei model was trained to distinguish all Brachyury-positive nuclei. The whole notochord model assigns bright Brachyury-positive nuclei and their inter-nuclear space to the foreground class, and the region outside the notochord, including weak Brachyury-positive cells in the tailbud, to the background class. Probability maps output by each model were further processed in Python. Probabilities were smoothened using a Gaussian filter with sigma=0.2, and binarized using a threshold of 0.35. Binarized images were further processed by identifying objects using connected component labelling, removing stray objects smaller than 5000 pixels, and filling holes in each cross-sectional plane. Finally, the notochord nuclei mask was further refined to exclude background and tailbud nuclei by multiplying it by the whole notochord mask.

The cross-sectional area and total volume of the notochord were calculated from the refined nuclear segmentation mask. For every cross-sectional plane of the notochord, the concave hull was computed to approximate the notochord boundary, enclosing the nuclear area and inter-nuclear space. The concave hull was defined by triangulating the pixel coordinates of notochord nuclei using Delaunay triangulation, retaining triangles whose edge lengths were less than 9 µm (approximate width of a cell), and merging the triangles into a single polygon. The area of the polygon was taken as the notochord cross-sectional area at every AP position along the notochord.

AP position was normalized such that the anterior edge equals 0 and the position of the posterior peak in cross-sectional area equals 1. For DTA embryos that lacked a clear posterior peak in cross-sectional area, the notochord posterior edge = 1. The mean and 95% confidence interval of the cross-sectional area were calculated for each developmental stage after AP position rescaling. To plot the cross-sectional area across absolute AP position, the rescaled AP position was multiplied by the mean notochord length at the respective developmental stage.

To estimate the number of cells in the notochord, the volume of all nuclei was divided by the mean volume of a single nucleus measured from a subset of at least 200 nuclei from 2-3 embryos for every stage using the nnInteractive plugin (Isensee et al., 2025) in the napari viewer for Python. Cell density was calculated by dividing the number of notochord nuclei by the total notochord volume. To obtain spatial profiles of cell density, volume quantifications were obtained in AP spatial bins of 1000µm starting from the posterior edge of the notochord.

To estimate the cell proliferation rate, first the number of phospho-Histone 3+ T/Bra+ cells was divided by the number of notochord cells quantified as described above. The fraction of diving cells was further divided by the duration of mitosis, which was assumed to be similar to that in the neural tube and equal to 28 minutes (Kicheva 2014). Spatial profiles of the proliferation rate were obtained by dividing the notochord into 3, 6 or 9 bins at E8.5, E9.5 and E10.5 respectively.

The number of apoptotic notochord cells was quantified by counting the number of cleaved caspase 3 (CC3) positive punctae in the Bra+ notochord in Imaris. This was divided by the total Bra+ cell number in the region of interest to calculate the apoptotic index.

To quantify the approximate AP extension rate of tissues external to the notochord, the following surrounding regions (SR) were defined: anterior region (A.N.) – between the anterior notochord edge and the anterior edge of the 1st somite, somites 1-9, and posterior region – from somite 10 to the posterior notochord edge. SR length was measured at the axial midline at distinct time points (Fig. 3I, S2G). The mean cumulative SR length (from the anterior edge of the notochord to the posterior boundary of every region) increased approximately linearly over developmental time as *SR*(*t*) = *SR*_0_ + *u_e_t*, where *u_e_* is the instantaneous velocity of the respective landmark. Using the cumulative SR length measurements, we determined *u_e_*of landmarks along the AP axis over developmental time (Fig. 4A). To estimate the spatial profile of the AP extension velocity of surrounding tissues, the velocity estimates for landmarks along the AP axis were fit to 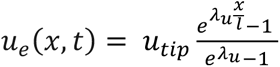 (Fig. 4B) at the initial time point, with *u_tip_* and *l*(*t*) fixed from a linear fit to total notochord length over time (Fig. 1D), and the linear fit for the cumulative SR length extrapolated to calculate the AP position of landmarks S16 and S22 at the initial time point, leaving the spatial shape parameter *λ_u_*(*t*) as the single free parameter at each stage.

To calculate the predicted growth of the notochord over time, the notochord was divided into regions as illustrated in Figure 2D. In each region, the expected number of cells generated by cell proliferation in 24 hours *N_t_*_24_ was calculated as *N_t_*_24_ = *N_t_*_0_ ⋅ *e^θ^*^⋅24^, where *N_t_*_0_ is the measured initial number of cells in the region and *θ* is the measured notochord proliferation rate.

To assess whether mitotic cells are preferentially located towards the edge of notochord fragments, we calculated the normalized distance of each mitotic (pH3-positive) cell to the nearest fragment edge. This yielded a value between 0 (mitotic cell located at the fragment edge) and 0.5 (mitotic cell located at the fragment midpoint). The normalized distances were pooled across all embryos and compared against a uniform distribution between 0 and 0.5 using a one-sample Kolmogorov-Smirnov test.

### Immunohistochemistry and microscopy of histological sections

Transverse cryosections were immunostained following a standard protocol. In brief, sections were defrosted in PBS at 42 °C, then incubated with primary antibodies in blocking buffer (1% BSA and 0.1% Triton-X100 in PBS) at 4°C for 2h. Sections were subsequently washed with 0.1% Triton-X100 in PBS, then incubated with secondary antibodies and DAPI (1:100) in blocking buffer at room temperature for 2 hours. Antibody dilutions: goat anti-Brachyury (1:250), rabbit anti-Laminin (1:1000), mouse anti-Nkx2.2 (1:25), rabbit anti-Olig2 (1:500), anti-goat FITC (1:250), anti-rabbit Alexa Fluor 647 (1:250)), anti-mouse Cy3 (1:250). Primary and secondary antibodies used are listed in Table M3. Transverse sections were imaged using a ZEISS inverted LSM800 confocal microscope, using a ZEISS Plan-Apochromat 20x/0.8 air objective or a ZEISS Plan-Apochromat 40x/1.2 water objective.

**Table M3.** Catalogue numbers for all antibodies used.

|  | **Company** | **Catalogue #** |
|---|---|---|
| **Primary antibodies** |  |  |
| Goat anti Brachyury/T | R&D Systems | AF2085 |
| Rabbit anti Laminin | Sigma | L9393 |
| Rat anti pH3 | Sigma | H9908 |
| Rabbit anti CC3 | Cell Signalling | #9661 |
| Mouse anti Nkx2.2 | DSHB | 74.5A5 (S) |
| Rabbit anti Olig2 | Millipore | AB9610 |
| **Secondary antibodies** |  |  |
| donkey anti goat IgG-FITC | Jackson ImmunoResearch | 705-096-147 |
| donkey anti rat IgG-Cy3 | Jackson ImmunoResearch | 712-166-153 |
| donkey anti rabbit IgG-Alexa Fluor 647 | Jackson ImmunoResearch | 711-606-152 |
| donkey anti mouse IgG-Cy3 | Jackson ImmunoResearch | 715-166-151 |

### Biophysical model

#### Governing Equations

We modeled the notochord as an axisymmetric rod of cross-sectional area *A*(*x*, *t*), with cells moving at velocity *u*(*x*, *t*). The conservation equation for a growing tissue takes the form:

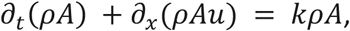

where *k*(*x*, *t*) denotes the local proliferation rate, and *ρ* is the cell density. As shown in Fig. 2A, the density *ρ* is spatially uniform and varies only with development time, i.e. *ρ* = *ρ*(*t*). Under this condition, the conservation equation simplifies to

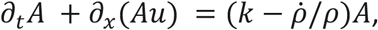

where *ρ̇* = d*ρ*/d*t*. Quantification of cell density over developmental time (Fig. 2A) revealed a linear increase. We capture this trend by setting *ρ* = *ρ*_0_(1 + *k_ρ_t*), with the rate constant *k_ρ_* = 0.0045/ℎ determined from the same data. This formulation makes explicit that local changes in cross-sectional area arise from three potential sources: gradients in cellular flux, cell proliferation, and changes in cell size or density.

On morphogenetic timescales, inertial effects are negligible and cell rearrangements justify treating the tissue as an active viscous material. The local force balance therefore considers three contributions: internal viscous stresses, friction with the surrounding basement membrane/ECM, and active migration forces.

For a viscous material with viscosity *η*, the axial stress in a rod undergoing uniaxial extension at rate *∂u*/*∂x* while growing at rate *k* is *σ* = 3*η*(*∂u*/*∂x* − *k*/3). The factor of 3 accounts for apparent shear thickening arising from geometric constraint and tissue incompressibility (Eggers and Dupont, 1994). The growth term −*k*/3 represents the contribution of growth to the spatial gradient of tissue velocity - omitting this growth term would lead to incorrect prediction of non-zero stress under homogeneous growth conditions.

The friction force per unit length between the notochord and the surrounding tissue is modelled as *ξA*(*u* − *u_e_)*, where *ξ* is the friction coefficient per unit area and *u_e_*(*x*, *t*) is the velocity field of the external tissue. Active cell migration generates a propulsive force *f_m_*.

Balancing these contributions yields:

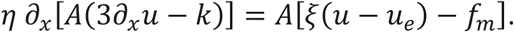

Non-dimensionalisation of the force balance reveals two key emergent parameters: a characteristic length 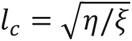 over which viscous and frictional forces equilibrate, and a propulsive velocity *u_m_* = *f_m_*/*ξ*, arising from active cell migration. The force balance equation becomes:

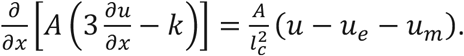

The anterior boundary is stationary in the laboratory frame, imposing: *u*(*x* = 0, *t*) = 0. At the posterior boundary *x* = *l*(*t*), the tissue velocity must match the boundary velocity: *u*(*x* = *l*, *t*) = d*l*/d*t*.

### Parameter estimation

Model parameters are constrained by experimental measurements as follows:

- The initial length *L*_0_ and tip velocity *u*_tip_ are directly determined from linear regression of length versus time data (Fig. 1D) as described in the main text.
- The initial area profile *A*(*x*, 0) is estimated from the measured area profile at E8.5 (Fig. 1E,F), which we parametrize as 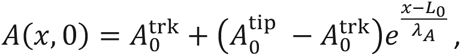 with 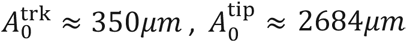 for wild-type embryos (Fig. 4B). The decay length was quantified as 124*μm* in Fig. 1I.
- To determine the relative increase rate of cell density, *ρ* = *ρ*_0_(1 + *k_ρ_t*) was fit to the data in Fig. 2A. This yields *k_ρ_* as the ratio of slope/intercept = 0.0045/h, which is the only parameter that enters the model, since *ρ*_0_ cancels out in *ρ̇*/*ρ*.
- The external velocity gradient *λ_u_* is a dimensionless parameter and is constrained by measurements of local elongation rates along the x-axis (Fig. 4A). In the model, we fit the elongation rate of somites and the presomitic mesoderm which yields *λ_u_* = 4.5 (Fig. 4B).
- The characteristic length *l_c_* characterizes the spatial extent over which viscous and frictional forces equilibrate. The force balance equation implies that the relative velocity between the notochord and surrounding tissues (quantified in Fig. 3G) should approximately follow 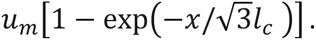 We estimate *l_c_* by fitting this relation to measured cell velocities obtained from live-imaging data along the x-axis (Fig. 3G), yielding *l_c_* ≈ 275*μm*.
- The active migration velocity *u_m_* is estimated as the plateau of velocity far from the anterior boundary in explant live imaging experiments (Fig. 3G). We find *u_m_* ≈ 9.4*μm*/ℎ.
- To consider a spatially modulated proliferation rate *k*(*x*, *t*) for a model without feedback, we implement 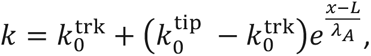 with 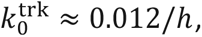 and 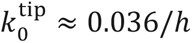 based on the experimental data in Fig. 2C.

The parameters used in the model are summarized in the Table M4:

**Table M4.**
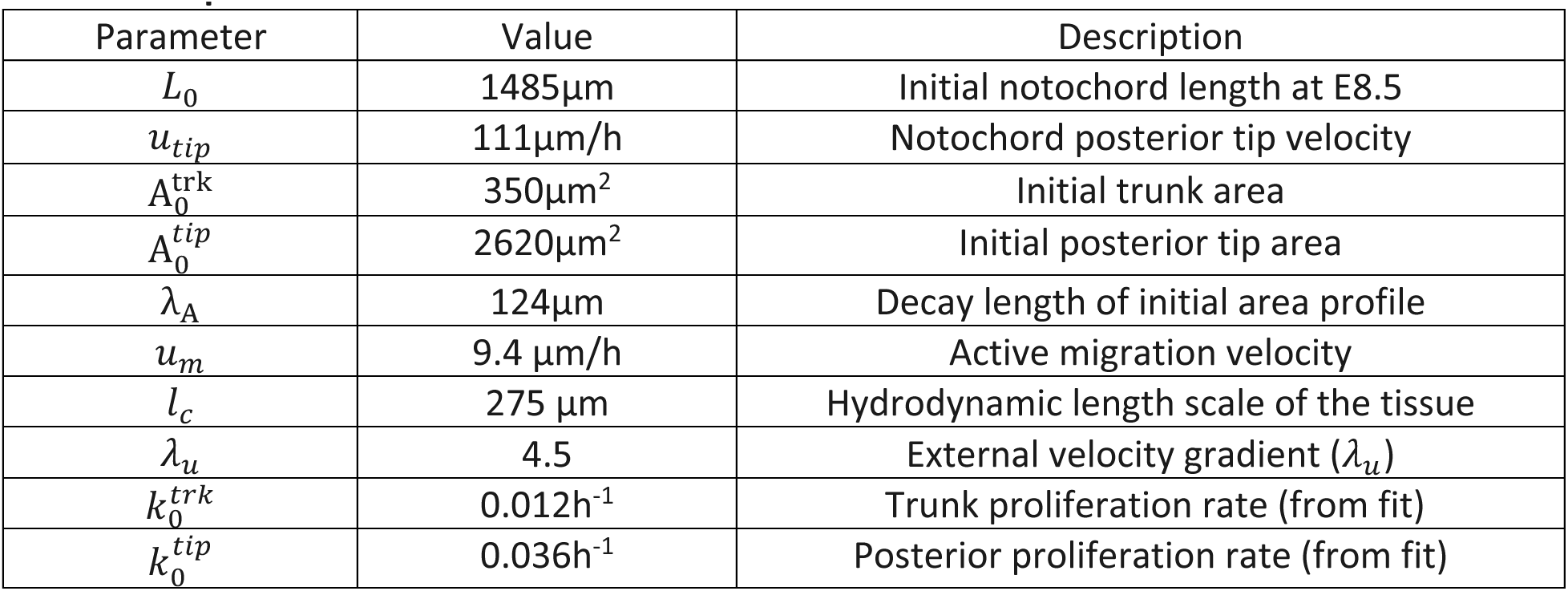
Core parameters of the model.

| Parameter | Value | Description |
|---|---|---|
| *L*<sub>0</sub> | 1485µm | Initial notochord length at E8.5 |
| *u*<sub>*tip*</sub> | 111µm/h | Notochord posterior tip velocity |
| A<sub>0</sub><sup>trk</sup> | 350µm<sup>2</sup> | Initial trunk area |
| A<sub>0</sub><sup>*tip*</sup> | 2620µm<sup>2</sup> | Initial posterior tip area |
| λ<sub>A</sub> | 124µm | Decay length of initial area profile |
| *u*<sub>*m*</sub> | 9.4 µm/h | Active migration velocity |
| *l*<sub>*c*</sub> | 275 µm | Hydrodynamic length scale of the tissue |
| *λ*<sub>*u*</sub> | 4.5 | External velocity gradient (*λ*<sub>*u*</sub>) |
| *k*<sub>0</sub><sup>*trk*</sup> | 0.012h<sup>-1</sup> | Trunk proliferation rate (from fit) |
| *k*<sub>0</sub><sup>*tip*</sup> | 0.036h<sup>-1</sup> | Posterior proliferation rate (from fit) |

#### Mechanosensitive growth feedback

To implement mechanically regulated proliferation, we assume that the growth rate is modulated according to:

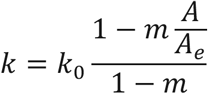

where *k*_0_ is the baseline growth rate in the absence of feedback, and *m* ∈ (0,1) is the feedback strength. *A_e_*(*x*) is the position-dependent target area, analogous to the carrying capacity in population dynamics, or to the homeostatic pressure in previous models of active matter (Basan et al., 2009; Hannezo et al., 2011; Podewitz et al., 2015; Recho et al., 2016). For an elastic ECM surrounding the notochord, at leading linear order this feedback arises from the resisting pressure exerted by the ECM on notochord cells. When *A* < *A_e_*, this pressure is reduced, permitting enhanced growth. In this view, the value of *A_e_* reflects the degree of mechanical/spatial confinement imposed on notochord cells in a specific position x by the surrounding ECM. The pre-factor 1/(1 − *m*) ensures that the proliferation rate is bounded to the baseline, i.e. *k* = *k*_0_ when *A* = *A_e_*.

The formulation above assumes a spatially uniform target area *A_e_*. However, the ECM surrounding the notochord has had more time to mature in the developmentally older anterior region compared to the posterior, suggesting that the degree of confinement varies along the AP axis and motivating a position- dependent *A_e_*. To incorporate this in the model, we allow:

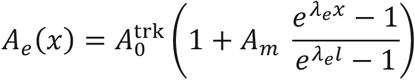

where 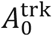 is the trunk target area, *A_m_* controls the amplitude of the posterior increase, and *λ_e_* determines the spatial decay of that target area. Larger values of *A_e_* in the posterior effectively weaken the feedback there, allowing larger deviations from the baseline area. We note that the model is robust to slight differences in the mathematical form of *A_e_*. The parameters used in the feedback formulation are found in Table M5.

**Table M5.**
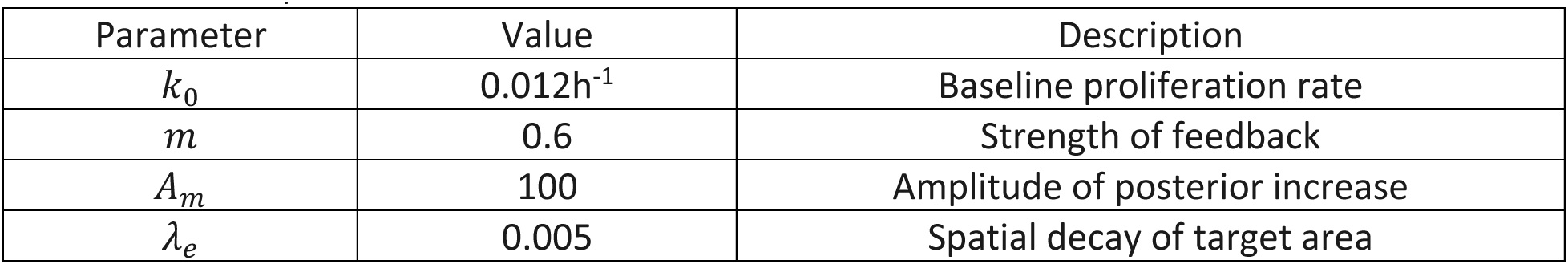
Feedback parameters used in the model.

### Implementation of perturbations in silico

i. **LPT mutants.** To capture the phenotypes of LPT mutants, which exhibit a reduced tissue elongation rate and a correspondingly lower tip velocity utip, we parameterize the model with utip = 79μm/ℎ by fitting the length data in Fig. 5B. The area profile was also modified for LPT mutants. Fitting the E8.5 data to 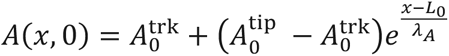 yielded 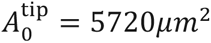 and λA = 137μm, while the initial trunk area 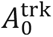 was kept as its wild-type value.
ii. **DTA mutants.** To model the transient cell death included in DTA mutants, we introduced a brief apoptotic window at the onset of E8.5, with duration ta = 6ℎ and apoptosis rate ka = 0.1/ℎ. Importantly, our qualitative conclusions are robust to variations in these parameters: DTA mutants consistently exhibit a reduced notochord cross-sectional area alongside an elevated cell proliferation rate.
iii. **Inhibition of cell migration.** We first performed a sensitivity analysis to examine the effect of cell migration on posterior area, where the “relative cell migration” denotes the value normalized to the wild-type level (Fig. 7A). Based on this analysis, we modeled inhibition of active notochord cell migration by reducing *u_m_* to 20% of its baseline level (Fig. 7B).

The parameters used in the perturbation implementation are found Table M6:

**Table M6.**
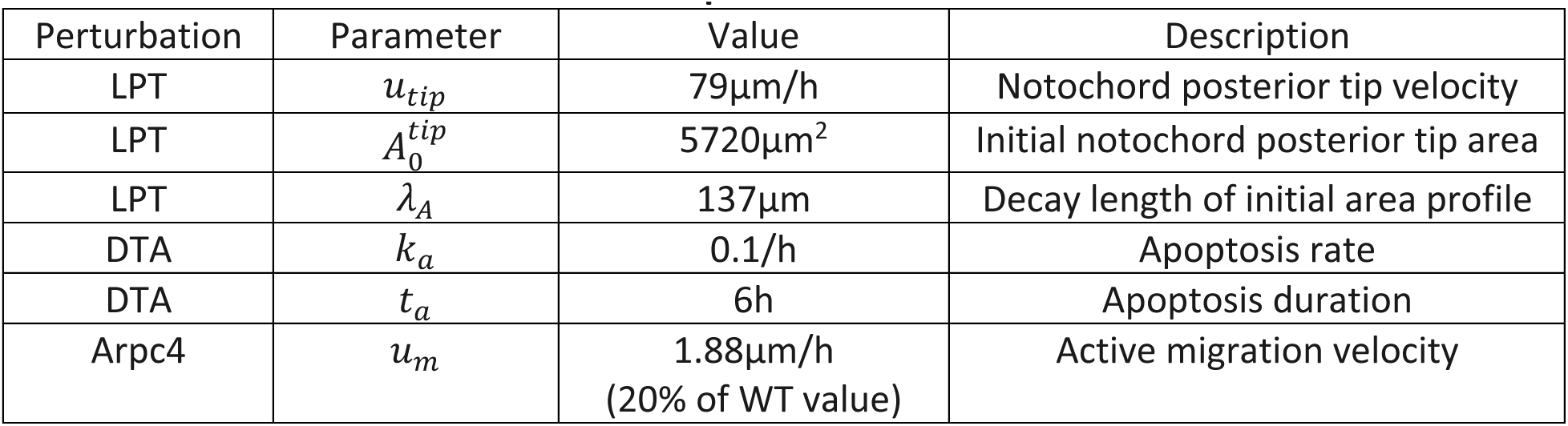
Parameter values used to simulate perturbed conditions.

| Perturbation | Parameter | Value | Description |
|---|---|---|---|
| LPT | *u*<sub>*tip*</sub> | 79µm/h | Notochord posterior tip velocity |
| LPT | *A*<sub>0</sub><sup>*tip*</sup> | 5720µm<sup>2</sup> | Initial notochord posterior tip area |
| LPT | *λ*<sub>*A*</sub> | 137µm | Decay length of initial area profile |
| DTA | *k*<sub>*a*</sub> | 0.1/h | Apoptosis rate |
| DTA | *t*<sub>*a*</sub> | 6h | Apoptosis duration |
| Arpc4 | *u*<sub>*m*</sub> | 1.88µm/h<br>(20% of WT value) | Active migration velocity |

#### Numerical Implementation

Our model forms a coupled system of partial differential equations that we solve numerically. We employ Comsol Multiphysics 5.4 to conduct the calculations.

## Supplementary Figures and Legends

**Figure S1:**
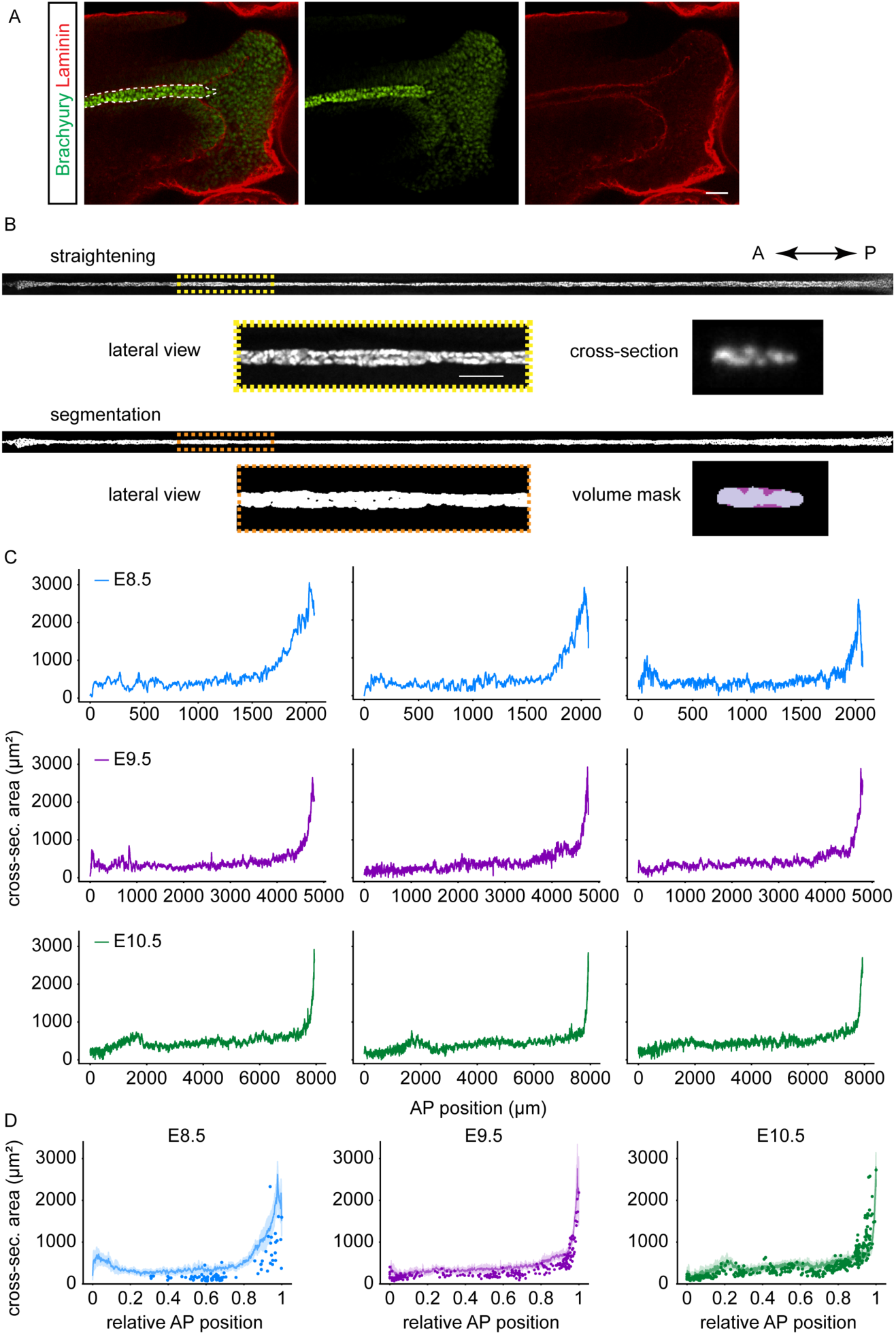
**(A)** The posterior notochord boundary is defined using strong Brachyury expression (white dashed line) throughout the study. Laminin (red) outlines the notochord and the weak Brachyury expression in the tailbud. Scale bar 50µm. **(B)** Notochord analysis from whole mounted cleared embryos. Top, lateral view of a digitally straightened notochord with zoomed region (yellow dashed rectangle) with scale bar 50µm, and cross-sectional view. Bottom, corresponding binary mask after segmentation. Zoomed region (orange). In the cross-sectional view, mask from the notochord nuclear segmentation (light purple). The sum of the light and dark purple regions shows the concave hull mask (notochord volume). **(C)** Example cross-sectional area profiles of individual embryos at the indicated stages. **(D)** Mean notochord cross-sectional area and 95% CI (shaded) from whole mount images, plotted together with notochord area quantified from serial cryosections (points) at the indicated stages.

**Figure S2:**
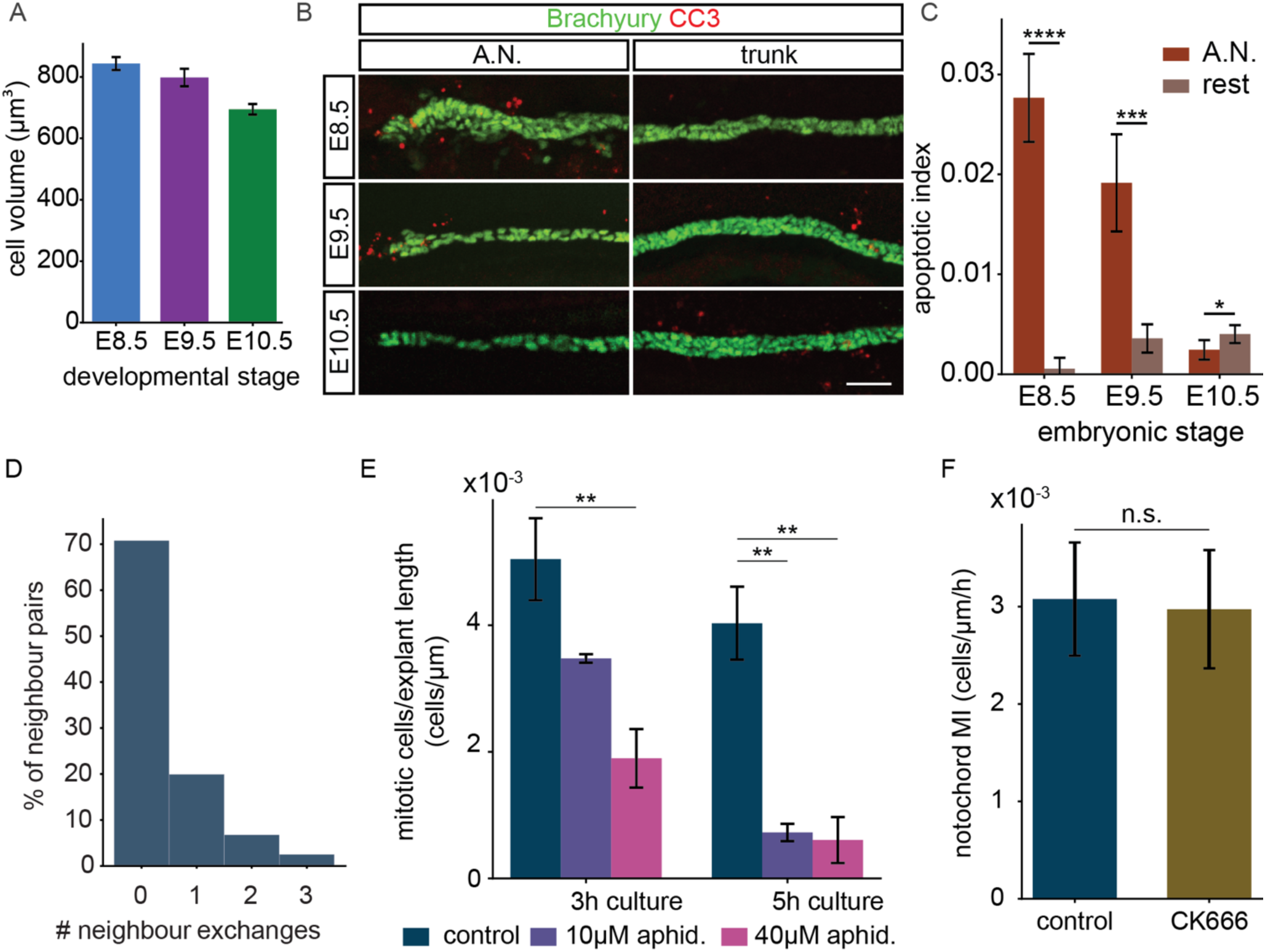
**(A)** Mean cell volume from E8.5 to E10.5. Error bars, SD. Sample size E8.5 n=12, E9.5 n=9, E10.5 n=6. **(B)** Immunostaining against cleaved caspase 3 (CC3) in the anterior (A.N.) and trunk regions from E8.5 to E10.5. **(C)** Quantification of the apoptotic index (CC3/Bra cells) in the anterior region and the rest of the notochord, from E8.5 to E10.5. Error bars, SD. T-test, p-value * p=0.028, *** p=0.00013, **** p<0.0001. Sample size E8.5 n=4, E9.5 n=5, E10.5 n=5. **(D)** Frequency of neighbor exchanges undertaken by pairs of neighboring cells in explant live imaging. Exchanges are defined as events in which cells get separated by more than a cell diameter (10 µm). **(E)** Quantification of the notochord mitotic index in explants treated with aphidicolin at the indicated concentration and culture duration. Error bars, SEM. ANOVA for 3h culture p=0.012, 5h culture p=0.0007. Asterisks, p-values from Tukey’s post-hoc HSD test, **p<0.01. Sample size DMSO n=4-5, 10µM aphidicolin n=2-3 for, 40µM aphidicolin n=4. **(F)** Notochord proliferation rate (number of mitotic cells per explant length per hour of live imaging) in explants treated with 800µM CK-666 for 6h. Error bars, SEM. T-test, p=0.91. Sample size n=11 for control, n=6 for CK-666. **(G)** AP length of regions surrounding the notochord over time. A.N. region anterior to first somite, S1-9=somites 1 to 9, P.N. region somite 10 to posterior notochord edge.

**Figure S3:**
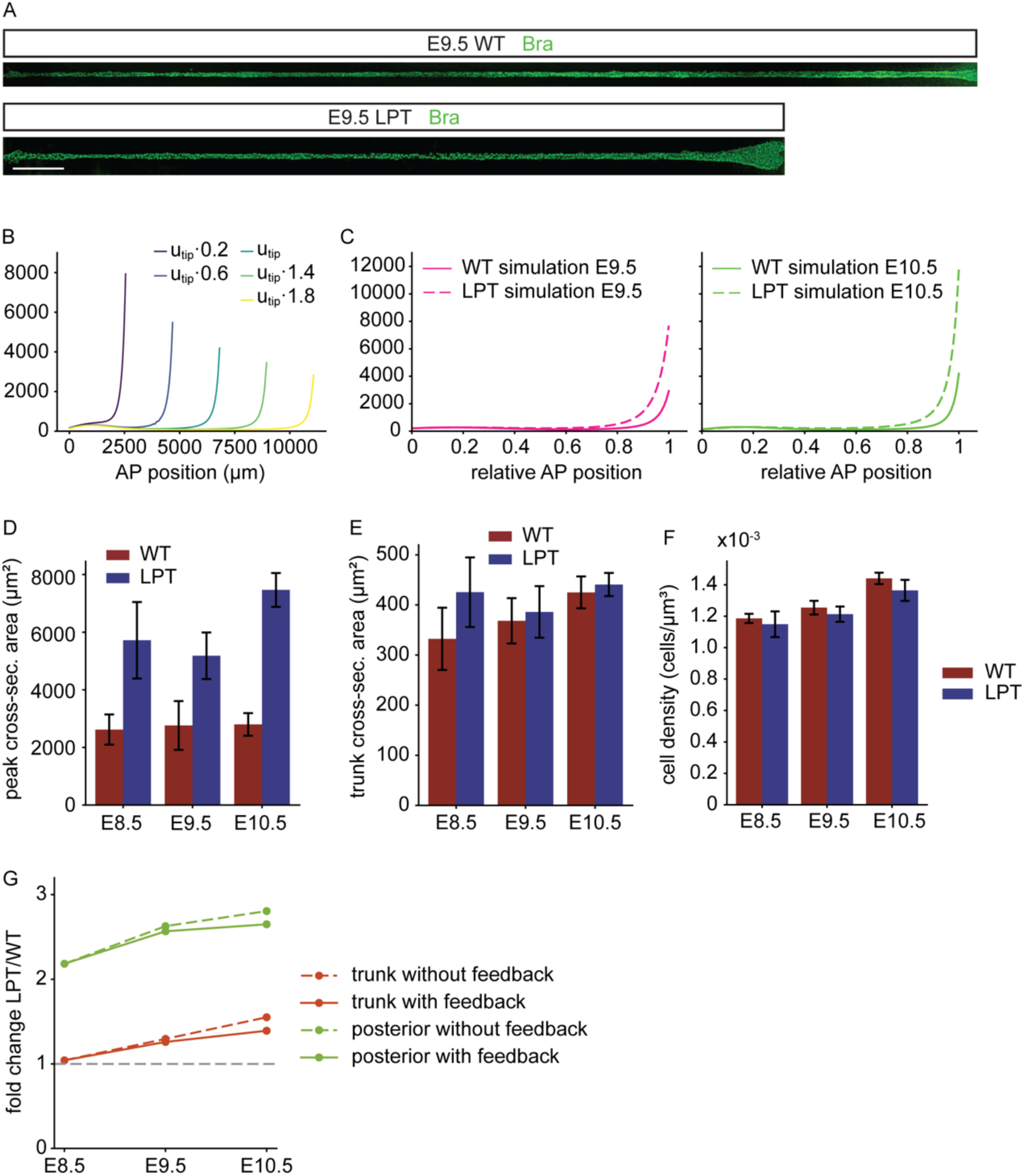
**(A)** Images of digitally straightened wildtype and homozygous LPT notochords at E9.5 of development immunostained for Bra. Scale bar, 200 µm. **(B)** Simulated cross-sectional area profiles for wildtype parameters and variable *u_tip_* for the model without feedback. **(C)** Cross-sectional area profiles resulting from the model without feedback (as in Fig. 4E) for the default wildtype parameters and decreased *u_tip_* condition, representing LPT mutants (Methods). **(D)** Quantification of the posterior peak cross-sectional area in wildtype and LPT embryos from E8.5 to E10.5. Error bars, SD. **(E)** Quantification of the trunk cross-sectional area in wildtype and LPT embryos from E8.5 to E10.5. Error bars SD. **(F)** Quantification of the notochord cell density in wildtype and LPT embryos from E8.5 to E10.5. Error bars, SD. Sample size in C, D, E as in Fig. 5C. **(G)** Ratio of the LPT to WT cross-sectional area of the whole trunk (red) or posterior peak (green) in the models with feedback (solid) or without feedback (dashed). Larger deviations from 1 indicate weaker robustness to the LPT perturbation.

**Figure S4:**
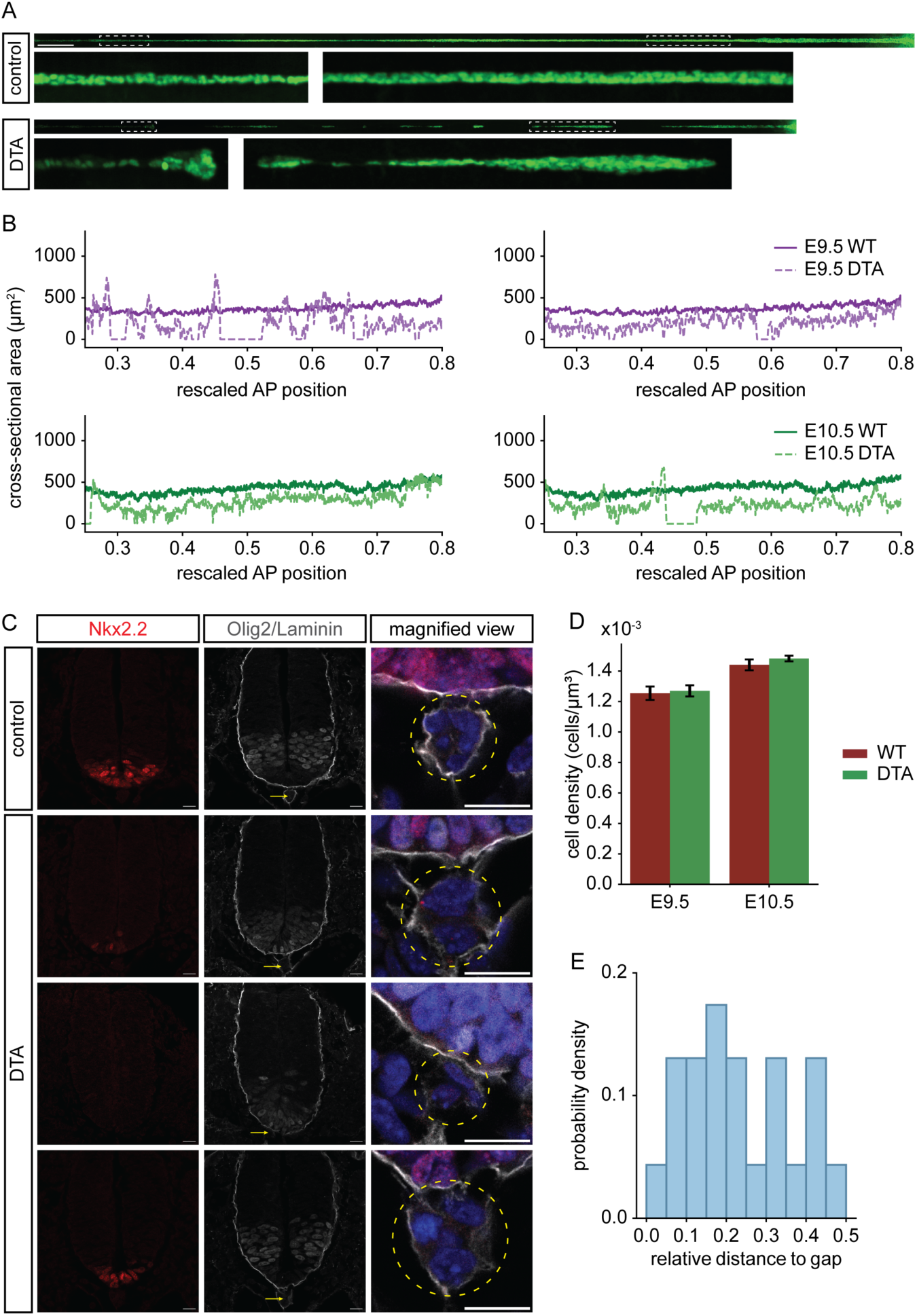
**(A)** Digitally straightened control and a DTA littermate notochord stained for Bra at E9.5. Dotted white boxes demarcate zoomed regions. Scale bar, 200µm. **(B)** Example cross-sectional area profiles of individual DTA embryos (dashed) in the trunk region, at E9.5 (purple) and E10.5 (green). Mean wildtype notochord cross-sectional area (solid) is shown for reference. **(C)** Transverse sections of the neural tube and notochord (arrow) of control and a DTA embryo at E9.5 immunostained for Nkx2.2 and Olig2. DTA sections from different AP positions in the same embryo display varying patterning defects. Third column, magnified view of the notochord (yellow circle) with DAPI (blue), Laminin/Olig2 (gray), Nkx2.2 (red). Scale bar, 15µm. **(D)** Quantifications of notochord cell density in WT and DTA embryos. Error bars, SD. Sample size E8.5 n=6, E10.5 n=8 **(E)** Probability density of the normalized AP distance of mitotic cells relative to the nearest fragment edge (gap). A value of 0 indicates the mitotic cell is located at the fragment edge, and 0.5 indicates it is located at the fragment midpoint. KS test comparison to a uniform distribution, p-value=0.83.

**Supplementary video M1**: Time-lapse imaging of a E9.5 mouse embryo trunk explant with sparse membrane-GFP labelled notochord cells (Shh-CreERT2::mT/mG) in control (DMSO) culture condition. The overlaid transmitted light channel shows neighbouring somites.

**Supplementary video M2:** Time-lapse imaging of a E9.5 mouse embryo trunk explant with sparse membrane-GFP labelled notochord cells (Shh-CreERT2::mT/mG) in 40µM aphidicolin culture condition. The overlaid transmitted light channel shows neighbouring somites.

**Supplementary video M3:** Time-lapse imaging of a E9.5 mouse embryo trunk explant with sparse membrane-GFP labelled notochord cells (Shh-CreERT2::mT/mG) in 800µM CK-666 culture condition. The overlaid transmitted light channel shows neighbouring somites.

